# Myosins generate contractile force and maintain organization in the cytokinetic contractile ring

**DOI:** 10.1101/2021.05.02.442363

**Authors:** Zachary McDargh, Shuyuan Wang, Harvey F. Chin, Sathish Thiyagarajan, Erdem Karatekin, Thomas D. Pollard, Ben O’Shaughnessy

**Affiliations:** Department of Chemical Engineering, Columbia University, New York, NY 10027, USA; Department of Physics, Columbia University, New York, NY 10027, USA; Department of Cellular and Molecular Physiology, Yale University, New Haven, CT 06520, USA; Nanobiology Institute, Yale University, West Haven, CT 06516, USA; Department of Molecular, Cellular, and Developmental Biology, Yale University, New Haven, CT 06520, USA; Department of Molecular Biophysics and Biochemistry, Yale University, New Haven, CT 06520, USA; Department of Cell Biology, Yale University, New Haven, CT 06520, USA

**Keywords:** cytokinesis, myosin II, actin, contractile ring tension, fission yeast, actomyosin contractility

## Abstract

During cytokinesis cells assemble an actomyosin contractile ring whose tension constricts and divides cells, but the ring tension was rarely measured. Actomyosin force generation is well understood for the regular sarcomeric architecture of striated muscle, but recent super-resolution studies of fission yeast contractile rings revealed organizational building blocks that are not sarcomeres but irregularly positioned plasma membrane-anchored protein complexes called nodes. Here we measured contractile ring tensions in fission yeast protoplast cells. The myosin II isoforms Myo2 and Myp2 generated the tension, with a ~2-fold greater contribution from Myo2. Simulations of a molecularly detailed ring model revealed a sliding node mechanism for tension, where nodes hosting tense actin filaments were pulled bidirectionally around the ring. Myo2 and Myp2 chaperoned self-assembling components into the ring organization, and anchored the ring against bridging instabilities. Thus, beyond force production, Myo2 and Myp2 are the principal organizers, bundlers and anchors of the contractile ring.

## Introduction

In the late 1960s it emerged that force-generating actomyosin contractile machinery is not confined to muscle, with Schroeder’s discovery of the contractile ring whose constriction divides cells during cytokinesis (Schroeder, 1968). With ~ 5 × 10^11^ daily sequences of ring assembly, maturation and constriction in humans (Ross, 1998), this machinery enables cell proliferation and the propagation of genetic material with extraordinary fidelity (Glotzer, 2017; Normand and King, 2010; Pollard and O’Shaughnessy, 2019). However, the relation between organization and force is far less clear for the contractile ring than for striated muscle, whose highly ordered organization is based on the sarcomere repeat unit. Sarcomeres contract when parallel actin filaments engage and slide past Myosin II thick filaments (Huxley and Hanson, 1954).

How tensile force emerges from the irregular actomyosin ring organization remains an open question. Following the discovery of actin filaments (Schroeder, 1972) and Myosin II (Fujiwara and Pollard, 1976) in the ring, the parallels with muscle led Schroeder and others to propose a muscle-like sliding filament mechanism (Fujiwara and Pollard, 1976; Mabuchi and Okuno, 1977; Schroeder, 1972). However, cytokinetic rings lack the ordered sarcomeric architecture of muscle (Laplante et al., 2016; Reymann et al., 2016).

In recent years super-resolution microscopies have probed the contractile ring organization in unprecedented nanoscale detail (Beach et al., 2014; Bellingham-Johnstun et al., 2021; Laplante et al., 2016). FPALM imaging showed that constricting fission yeast contractile rings, uniquely well characterized for their components and amounts (Courtemanche et al., 2016; Friend et al., 2018; Wu and Pollard, 2005), have an organization based on protein complexes called nodes (Laplante et al., 2016). This finding suggests a major shift from the sarcomeric paradigm of muscle. Each node includes ~ 8 myosin II Myo2 dimers with plasma membrane-anchored tails, and the actin filament nucleator formin Cdc12, so actin filaments presumably emanate from nodes. With ~ 200 nodes distributed irregularly around the ring at constriction onset, the node is the building block of the contractile ring, not the sarcomere. How a disordered node-based organization generates contractile ring tension is not established.

A major obstacle to finding the relationship between organization and tension production has been the near absence of ring tension measurements. Classic studies reported net forces ~ 10-60 nN in the furrows of echinoderm eggs (Hiramoto, 1970; Rappaport, 1967, 1977; Yoneda and Dan, 1972), but these include force from contiguous actomyosin cortex, or unverified assumptions were invoked. More recently, we measured ~ 400 pN tensions in contractile rings of fission yeast protoplasts, live cells with the cell wall removed (Stachowiak et al., 2014). Given the paucity of ring tension data, constriction rates are typically used as a proxy (Laplante et al., 2015; Piekny and Mains, 2002).

Actomyosin organizations typically use several myosin II isoforms with complementary roles. The fission yeast contractile ring has a second isoform, the unconventional myosin II Myp2, not belonging to the nodes and likely unanchored from the membrane (Bezanilla et al., 1997). The respective roles of Myo2 and Myp2 are not known. Animal cells express multiple Myosin II isoforms (NMIIA,B and C), with the more dynamic NMIIA mediating rapid contraction, while NMIIB promotes isometric tension (Shutova and Svitkina, 2018). In contractile rings NMIIA and NMIIB assemble into stacked minifilaments (Beach et al., 2014) and furrow ingression is slower in NMIIA-knockout cells (Yamamoto et al., 2019).

Here we study the tension mechanism in the node-based organization of the fission yeast cytokinetic ring. Measurements of ring tension in live cells and mutants of the myosin II isoforms show that both isoforms contribute, with a ~ 2-fold greater contribution from Myo2. In simulations implementing the experimentally measured node organization, myosin generated tension by pulling node-attached actin filaments, explaining the observed bidirectional node motions around the ring (Laplante et al., 2016). Incoming components continuously self-assembled into the ring organization, chaperoned by Myo2 and Myp2 which maintain a dense actomyosin core and a dilute outer corona, respectively. Myo2 is the principal anchoring agent opposing bridging instabilities, the major threat to contractile ring integrity, and simulations reproduced bridging seen experimentally in the *myo2-E1* mutant with weak actin binding (Lord and Pollard, 2004). Thus, beyond exerting force that makes ring tension, Myo2 and Myp2 have complimentary roles as the principal organizers, bundlers and anchoring agents of the contractile ring.

## Results

### Method to measure contractile ring tension in fission yeast

Past efforts to measure contractile ring tension in animal cells were complicated by interference from the actomyosin cortex. Here we used fission yeast protoplasts, living cells with enzymatically digested cell walls (Mishra et al., 2012). As fission yeast lacks an actin cortex, contractile rings in protoplasts are isolated so ring tension is opposed only by plasma membrane tension (Stachowiak et al., 2014).

The ~ 5-7 μm diameter spherical protoplasts were osmotically stabilized by 0.8 M sorbitol, and <5% had contractile rings which slid along the membrane as they constricted, their tension furrowing the membrane (Figure S1). Micropipet Aspiration measured the membrane tension in one lobe, *σ*_1_, and Laplace’s law yielded the tension in the unaspirated lobe, *σ*_2_ = *σ*_1_ *R*_2_/*R*_1_, where *R*_1_, *R*_2_ are the lobe radii (Figure 1B and STAR Methods). The ring tension *T* follows from a force balance, *T*/*R*_ring_ = *σ*_1_*cosθ*_1_ + *σ*_2_*cosθ*_2_, where *R*_ring_ is the ring radius and *θ*_1_ and *θ*_2_the furrow angles. Angles and lobe radii were measured from images of the lobes with fluorescently labeled membrane. This improves our earlier method where membrane tension measured in interphase cells was used in the force balance in mitotic cells (Stachowiak et al., 2014). Membrane tensions were far lower in cells without rings (Figure S1A), so the previous method underestimated ring tensions.

**Figure 1.**
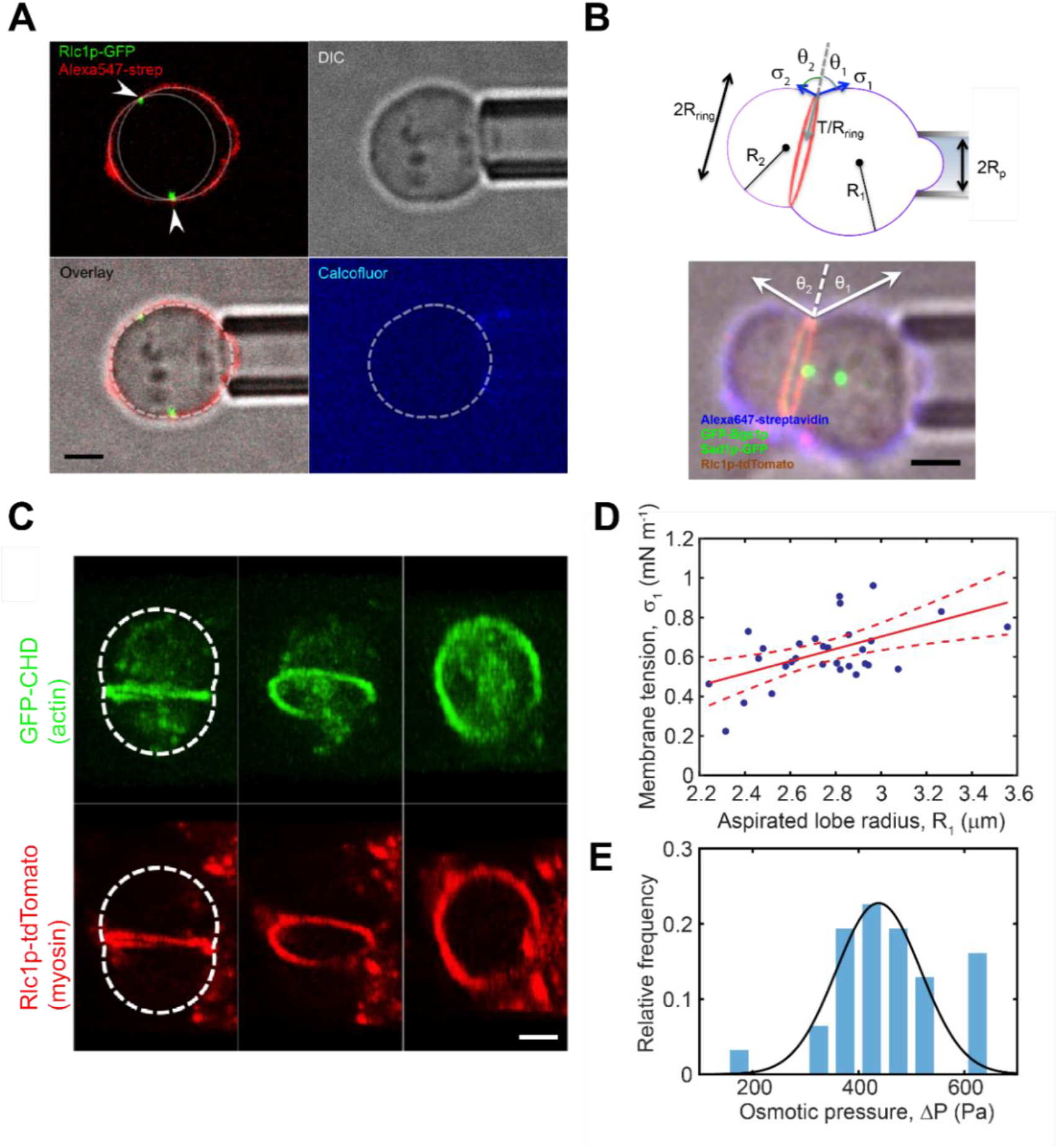
Method to measure cytokinetic ring tension in fission yeast protoplasts. **(A)** A protoplast cell is held by an aspirating pipette and furrowed by a cytokinetic ring. The ring is marked by myosin II light chain (Rlc1p-3GFP), the plasma membrane by Alexa555-streptavidin (red). The furrow geometry (arrowheads) was measured by fitting circles to the flanking lobes. Absence of calcofluor white staining indicates minimal regrowth of cell wall (lower right). Scale bars, 2 μm. **(B)** Top, geometry for ring tension measurements. Bottom, fluorescence and bright field overlay of an aspirated protoplast with labeled spindle pole bodies (Sad1p-GFP), myosin and Bgs1p in cytokinetic rings (Rlc1p-tdTomato, GFP-Bgs1p), and plasma membrane (Alexa647-streptavidin). **(C)** Confocal micrographs of a furrowed protoplast. Actin and myosin II colocalize in the cytokinetic ring, and actin is visible in the cell interior, but actomyosin cortex is absent. Images from left to right show 3D reconstructions consecutively rotated 40°. Dashed white line, cell boundary. **(D)** Membrane tensions of furrowed protoplasts versus aspirated lobe radius, R_1_ (*n* = 31). An approximately linear relation holds, with best fit slope 0.31 ± 0.09 mN/m/μm (red line, F test, *p <* 0.01, and Pearson correlation coefficient, *r* = 0.52; dashed curves, 95% confidence interval). **(E)** Histogram of osmotic pressures in furrowed protoplasts of (D), obtained from Laplace’s Law (Δ*P* = 2*σ*_1_/*R*_1_). Mean pressure is 440±30 Pa (mean ± SD), or osmolarity ~ 0.2 mM. Solid curve: best fit Gaussian.

Corroborating the method’s assumption of a spatially uniform plasma membrane tension, the lobes flanking ring-induced furrows were almost perfect spherical caps. Membrane tension scaled approximately as the aspirated lobe radius (*p* < 0.01, Figure 1D), indicating consistent osmotic pressures among cells (*ΔP* = 440 ± 30 Pa, mean ± SD, *n =* 31 cells, Figure 1E).

### Contractile ring tension increases from ~400 to ~800 pN as the ring constricts

We first used the method to measure the evolution of contractile ring tension as rings constricted. In measurements on 31 protoplast cells ring tensions increased as constriction progressed, from ~ 400 to ~ 800 pN (Figure 2) with a mean of 640 ± 290 pN (SD, *n* = 31).

**Figure 2.**
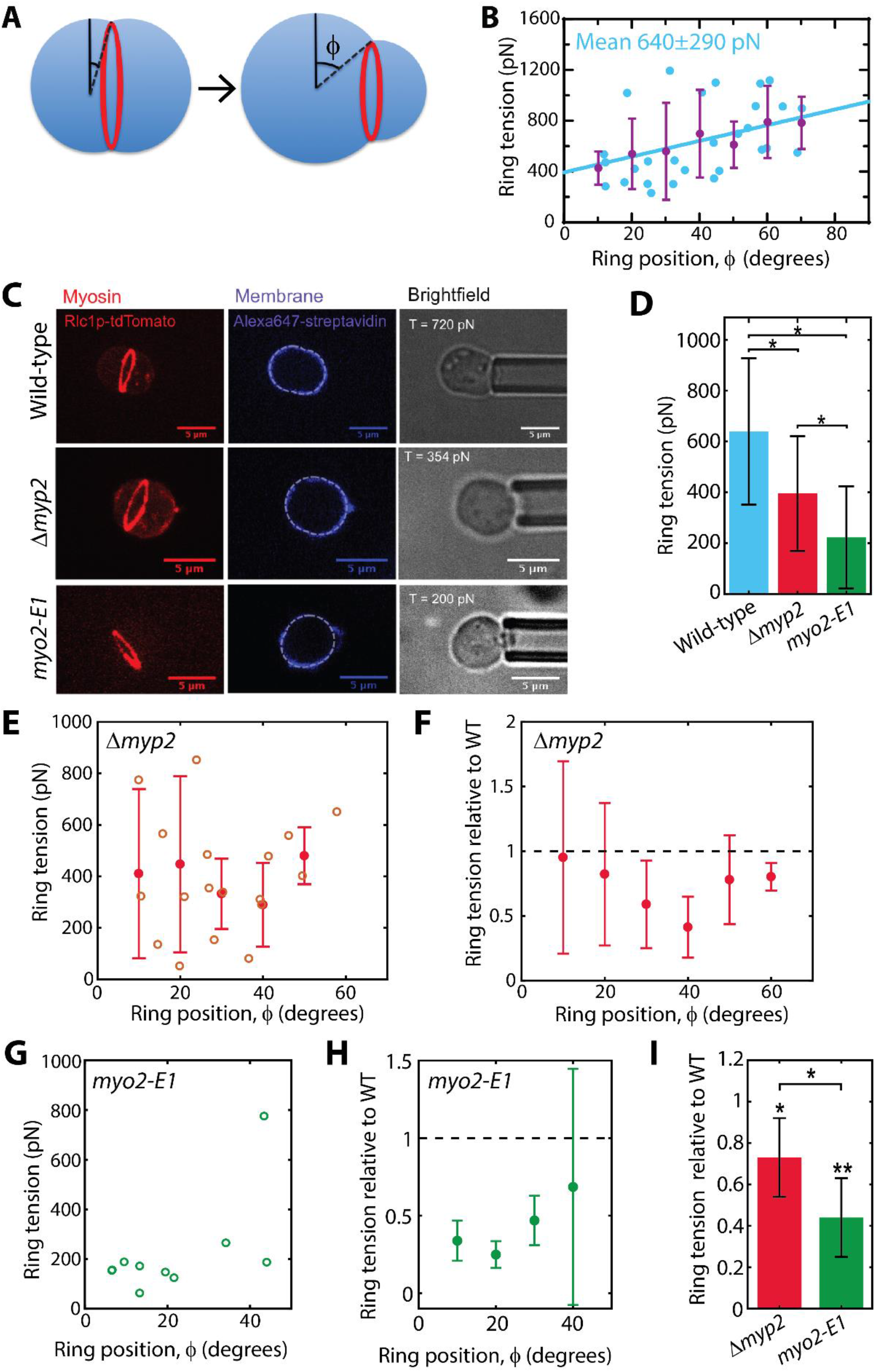
Cytokinetic ring tension increases as the ring constricts and is accounted for by the myosin II isoforms Myo2 and Myp2. **(A)** Schematic of a constricting ring in a protoplast, located by the angle *ϕ* between the ring and large lobe equatorial planes. As the ring constricts it slides along the inside of the protoplast membrane. **(B)** Top, ring tensions versus ring location *ϕ* (n=31). Ring tension increases as the ring constricts (Pearson coefficient r = 0.40; solid line, best linear fit). Purple, binned tensions (bin width, 10°). Best linear fit (solid line), 58 ± 10 pN per 10 degrees (*p* = 0.0023, F test). Error bars, SD. See Figure S2 for tensions versus time for individual constricting rings. **(C)** Confocal fluorescence and bright field images of WT, *Δmyp2* and *myo2-E1* protoplasts aspirated to measure the tension of a contractile ring (left column). Each row shows images of the same cell. Plasma membrane, dashed lines (middle column). Measured ring tension indicated (right column). Table S2 lists the *S. pombe* strains used in this study. **(D)** Ring tensions averaged over all ring positions for WT, *Δmyp2* and *myo2-E1* protoplasts. Error bars, SD. **(E)** Ring tensions in *Δmyp2* protoplasts (open circles, *n=*18), and binned tension values (closed circles, bin width 10°) versus ring location *ϕ*. Error bars, SD. **(F)** *Δmyp2* tension relative to WT tension versus ring location. For each bin of ring locations *ϕ* (bin width, 10°) the ratio of the binned *Δmyp2* tension in **(C)** to the binned WT tension in Figure 2B is plotted. Error bars, SEM. **(G)** Ring tensions in *myo2-E1* protoplasts versus ring location *ϕ* (*n* = 10). **(H)** *myo2-E1* tension relative to WT tension versus ring location. For each bin of ring locations the ratio of the binned *myo2-E1* tension from (E) to the corresponding binned WT tension in Figure 2B is plotted. Error bars, SEM. **(I)** Mean tension ratios for *myo2-E1* and *Δmyp2* protoplasts shown in **(D)** and **(F)**, averaged over angles *ϕ* (mean ± SD). \**p<*0.05, **p<0.01, Student’s t test.

Each tension in Figure 2B refers to a different ring at a different stage of constriction. We also tracked successive tensions in five rings as they constricted. The tensions increased at ≤ 5% per min (Figure S2). Given a ~ 50 min constriction time in protoplasts (Stachowiak et al., 2014), these rates are consistent with the single snapshots, for which tensions varied ~ 2-fold over the full constriction range.

### Contractile ring tension is ~30-40% lower in cells lacking Myp2p

We measured the tension contributions of the two myosin II isoforms, Myo2 and Myp2, using protoplasts of cells with mutations of each myosin gene (Figure 2C). In protoplasts generated from *Δmyp2* cells lacking unconventional myosin II Myp2, the mean ring tension in 18 cells with rings at various stages of constriction was 400 ± 230 pN (mean ± SD), ~ 38% below the wild-type (WT) mean of 640 pN (Figure 2D).

Compared at the same degree of constriction, the *Δmyp2* ring tensions were ~ 5% – 60% reduced from WT (Figures 2E & 2F). Averaged over all polar angles *ϕ* (0^0^ < *ϕ* < 60^0^), the reduction factor was 0.73 ± 0.19 (mean ± SD, *n* = 5 bins), or a mean ~ 27% tension loss (Figure 2I). These results suggest Myp2 contributes ~ 30-40 % of the total WT ring tension.

### Contractile ring tension is ~ 60% lower in the absence of Myo2 ATPase activity

The temperature-sensitive *myo2-E1* mutation (Balasubramanian et al., 1998) lacks ATPase activity (Lord and Pollard, 2004). The mean ring tension in 10 *myo2-E1* protoplasts was 220 ± 200 pN (mean ± SD), ~ 66% below the WT tension (Figure 2D). Rings constricted through a smaller range of angles (0^0^ < *ϕ* < 45^0^) than WT cells. Compared at the same degrees of constriction, tensions in *myo2-E1* protoplasts were ~ 44% ± 19% of WT values (mean ± SD, *n* = 4 bins) (Figures 3E, 3F & 3G).

**Figure 3.**
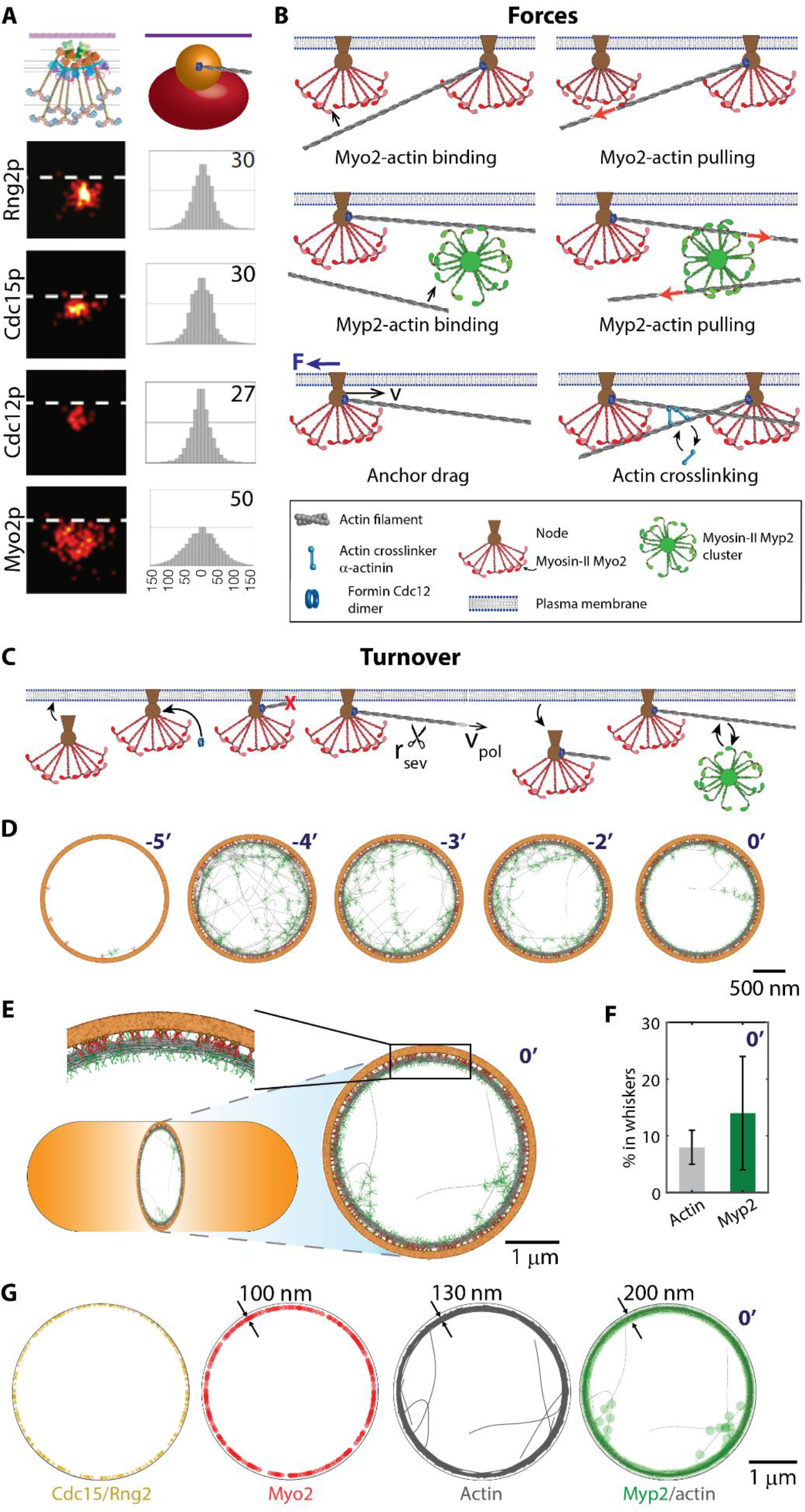
Mathematical model of the fission yeast contractile ring predicts self-assembly of components into a functional ring. Color code used in this and subsequent figures: Myo2 (red); Myp2 (green); Actin (grey). **(A)** The model implements the experimental node organization in constricting rings, including node stoichiometries and spatial distributions. Lower 4 rows, FPALM images of node components (left) and radial density distributions (right; radii with 75% of FPALM localizations in nm; plasma membrane, dashed white line). Top row, schematic of membrane-anchored node organization (left) and coarse-grained model representation of Myo2 heads (red) and Rng2 and Cdc15 (brown). Formin Cdc12 dimers bind the node 40 nm from the membrane. FPALM images, density distributions and node schematic reproduced with permission from (Laplante et al., 2016). **(B)** Forces in the model. **(C)** Turnover. Nodes bind and dissociate from the plasma membrane. Formin Cdc12 dimers bind nodes and grow randomly oriented actin filaments at rate *ν*_pol_. Cofilin stochastically severs filaments. Myp2 clusters bind/unbind filaments. **(D)** Starting from a bare membrane, ring components self-assemble into a functional actomyosin bundle within 5 mins (see Figure S6). Myo2 (red) and Myp2 (green) are rendered with explicit molecular detail for clarity. **(E)** Simulated ring at constriction onset, following 3 min of equilibration from a random initial condition. **(F)** Fraction of actin and Myp2 in whiskers at constriction onset (mean ± SD, *n* = 60 rings). **(G)** Spatial distributions of components in the steady state ring of (D).

Thus, Myo2 accounts for ~ 60% of the total WT ring tension. Together with the Myp2 contribution, the two myosin II isoforms appear to generate most or all contractile ring tension.

### Molecularly detailed simulation of the *S. pombe* cytokinetic ring

We built a molecularly explicit mathematical model tightly constrained by experiment (Figure 3A-C, Table S1, STAR Methods) to study how tension emerges from the node-based organization in the fission yeast ring. Our previous 2D fixed ring length model featured independently membrane-anchored formin Cdc12 and Myo2 (Stachowiak et al., 2014). The 3D model developed here implements the measured node-based organization, incorporates Myp2 and evolves the length of the ring as it constricts. For details of the model, simulation method and parameters see STAR Methods and Table S1.

#### Node organization

The simulations implement the local organization revealed by super resolution microscopy, which showed Myo2, formin Cdc12, the IQGAP Rng2p and the F-BAR protein Cdc15p colocalizing in membrane-anchored nodes (Laplante et al., 2016) (Figure 3A). In simulations: (i) Near the membrane, a sphere represents Cdc15p and IQGAP Rng2p, with size and location matching the experimental 70 nm wide Cdc15p distribution. (ii) Formin Cdc12 dimers nucleate and grow actin filaments, binding the node 44 nm from the membrane. (iii) An ellipsoid with the experimental 132 × 102 × 102 nm dimensions located 94 nm from the membrane represents the heads of the ~ 8 Myo2 dimers.

#### Other components

Semiflexible actin filaments are represented by chains of rods with bending stiffness corresponding to the ~ 10 μm persistence length (Ott et al., 1993). The unconventional Myosin II, Myp2, forms puncta in constricting rings suggestive of clustering (Takaine et al., 2015), does not belong to nodes and is likely not directly anchored to the plasma membrane (see Discussion). We simulated Myp2 as unanchored 200 nm clusters of 16 molecules (Figure 3B), the values best reproducing experimental ring tensions and organization (Figure S3). The model did not include type V myosin Myo51, which dissociates before constriction completion (Laplante et al., 2015), or formin For3 since cytokinesis appears normal in *Δfor3* cells, including normal actin levels in rings (Feierbach and Chang, 2001).

#### Actin-myosin forces

Filaments intersecting Myo2 or Myp2 clusters bind and are pulled parallel to the filament (Figure 3B). As the filament unbinding forces are unknown, we scanned values and selected those best reproducing the experimental radial Myo2-Myp2 separation and actin bundle thickness (Figure S4). We used linear force-velocity relations for Myo2 and Myp2, with stall forces per cluster, 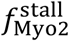 and 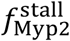, best reproducing our experimental ring tensions (Figure S5), and with the experimentally measured load-free Myo2 velocity 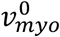 or the estimated 25 myosin II heads per actin filament 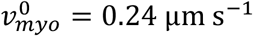 (Stark et al., 2010). The same load-free velocity was used for Myp2, as Myo2 and Myp2 heads appear similar (Bezanilla and Pollard, 2000). The cluster force is evenly distributed over all bound filaments.

#### Other forces

Drag forces resist node membrane anchor motion parallel to the membrane, with a drag coefficient chosen to reproduce the experimental node velocity, 22±10 nm s^−1^ (Laplante et al., 2016). Node-node, Myp2-Myp2, node-Myp2 and actin-actin steric forces oppose overlapping of clusters or filaments (thus filaments cannot cross). Actin filaments are dynamically crosslinked by Ain1p α-actinin dimers, the ring’s most abundant passive crosslinker (~ 250 at constriction onset) (Wu and Pollard, 2005).

#### Amounts of ring components and turnover

The component amounts follow the experimental values throughout constriction (Courtemanche et al., 2016; Wu and Pollard, 2005), e.g. at onset simulated rings contain 3300 Myo2 molecules and 230 formin dimers in ~210 nodes, 2300 Myp2 molecules in ~140 clusters, and 230 actin filaments of total length ~580 μm. Nodes stochastically bind a 0.2 μm wide zone, representing the ingrowing septum edge (Cortes et al., 2007), and unbind after a mean of 41 s consistent with reported component dissociation times (see STAR Methods). On binding a node, a formin nucleates and grows an actin filament that is stochastically shortened by cofilin-mediated severing (Elam et al., 2013). Results were insensitive to the nucleated filament orientation distribution. Myp2 clusters bind actin filaments with turnover time 46 s (Takaine et al., 2015). Rates of binding, actin polymerization and actin filament severing are set by demanding simulations reproduce the experimental densities of Myo2, Myp2 and formin and mean actin length, all evolving as constriction progresses.

#### Running the simulation

A dynamic boundary condition represented the ingrowing septum whose diameter, initially 3.7 μm, decreased at 2.4 nm s^−1^ (Thiyagarajan et al., 2015) for a total constriction time ~ 25 min. The width of the contractile ring was self-determined by the simulation.

### Ring components self-organize into a functional contractile ring

Remarkably, in simulations of the model the incoming components self-organized into a contractile ring whose organization and tension reproduced experiment, independently of initial conditions (Figures 3-5). Even with an initially empty membrane, the stochastically incoming Myo2-containing nodes, Myp2 clusters, formins and freshly-nucleated actin filaments assembled without supervision into the same bundled actomyosin organization with the same tension, (Figures 3D & S6).

Typical simulations used an initial condition with components randomly positioned in a 200 nm wide circular band, STAR methods. Before initiating constriction, the simulation was run for 3 mins until the ring self-organized. This dynamic organization maintained itself throughout constriction and reproduced many features measured in live cells (Figures 4E-G).

**Figure 4.**
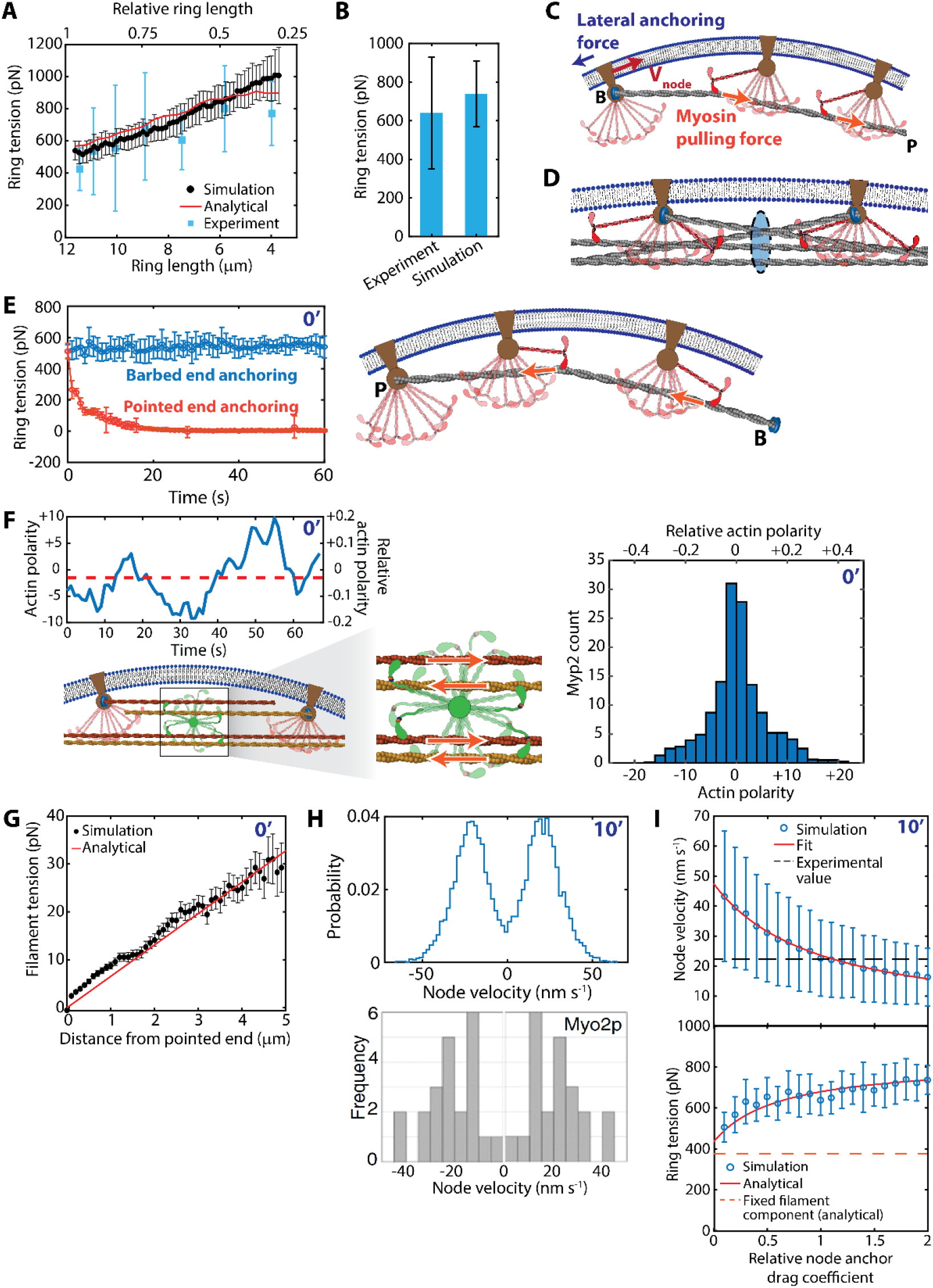
Sliding node ring tension mechanism. **(A)** Simulated ring tensions WT (black, mean ± SD, n = 60 simulations, parameters as in Table S1). WT experimental tensions (blue, data from Figure 2B, compared at same length relative to initial length). Approximate analytical expression, Equation (1) (red curve). **(B)** Mean WT tensions from experiment (n=31, Figure 2B) and simulations (n = 60 simulations). Error bars are SD. **(C)** Sliding node ring tension mechanism. On average a node anchors the barbed end of one actin filament to the membrane. Pulled by myosin, filament tension results since the node-membrane anchor drag force large enough that the node velocity *ν*_node_ is well below the load-free myosin II velocity. **(D)** The ring tension sums tensions of all filaments passing through a given ring cross-section. Nodes with oppositely oriented actin filaments are drawn together. **(E)** Pointed end anchoring fails to generate tension. Tension was lost within ~30 seconds (left, *n* = 20, error bars are SD) as filaments are compressed or buckled by myosin II (right). **(F)** Unanchored Myp2 clusters contribute to tension by pulling on equal numbers of oppositely oriented filaments. Net filament polarity bound to a Myp2 cluster is small and fluctuates in time (top left). Net polarity histogram for all Myp2 clusters in a contractile ring (n=10 rings at constriction onset, right). Relative polarities are normalized by the maximum value (all filaments having same polarity). **(G)** Mean tension profile along one actin filament, constriction onset (mean ± SEM, n = 174 filaments). Red line: expected linear profile for uniform myosin II binding (see main text and STAR Methods). **(H)** Nodes move bidirectionally around the ring. Top: bimodal distribution of node velocities in simulated rings (*n* = 60 simulations, ~120 nodes per ring, 500-600 s after constriction onset). Nodes were tracked for 40s. Bottom: Myo2 velocity distribution measured in cells, reproduced from ref. (Laplante et al., 2016). **(I)** Normal ring tension requires firm anchoring of actin filament barbed ends to the plasma membrane. Lowering node anchor drag coefficient decreases simulated ring tensions, approaching the drag-independent fixed filament component (n=10 rings per data point, 10 min after constriction onset, error bars represent SD). The tension data are well described by an analytical model (red curve, STAR Methods). Anchor drags are relative to the WT simulated value (see main text).

The actin filaments aligned into a circular bundle of mean thickness 131±4 nm (mean ±SD, *n* = 60 rings at constriction onset) (Figure 3G), within the reported range from FPALM and cryoET, ~100 – 175 nm (Laplante et al., 2016; McDonald et al., 2017; Swulius et al., 2018). A fraction ~9 % of actin belonged to “whiskers” emanating from the bundle (Figures 3E and 3F) as observed in live cells (Vavylonis et al., 2008). A 100 nm thick Myo2 ring overlapped a concentric 200 nm thick inner Myp2 ring, both associated with the actin bundle (Figure 3G), quantitatively consistent with experiment (Laplante et al., 2015; McDonald et al., 2017).

### Ring tension is generated by anchoring of actin barbed ends to the membrane

Simulated contractile rings produced tensions close to the values we measured in live cells, including the tension increase throughout constriction (Figures 2 and 4A). Compared to the experimental mean ~ 640 pN, the simulated mean was ~ 440 ± 140 pN (Figure 4B), using best fit stall forces of 1.75 pN (Myo2) and 1 pN (Myp2) from a systematic scan. These lie in the range reported for smooth and skeletal muscle myosin II, 0.7 – 2.1 pN (Ishijima et al., 1996; Tyska et al., 1999).

The tension mechanism relied on firm lateral anchoring of actin filament barbed ends to the plasma membrane (Stachowiak et al., 2014; Wang and O’Shaughnessy, 2019) via formins belonging to nodes (Alonso-Matilla et al., 2019; Thiyagarajan et al., 2017) (Figure 3A, B). Since myosin II tries to move toward barbed ends, this guaranteed filament tension (Figure 4C). Other hypothetical anchoring schemes generated less tension, or compression (Figure 4E). Both Myo2 and Myp2 contributed to tension; despite Myp2 being unanchored it also contributed, by pulling equally in both directions on the actin bundle of approximately zero net polarity (Stachowiak et al., 2012), Figure 4F.

This mechanism was apparent from the tension in individual actin filaments, *T*_fil_(*x*), which obeyed the relation for a barbed-end-anchored filament pulled by a density *C*_bound_ of myosin II along its length, 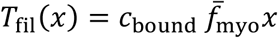 (*x* is distance from the pointed end and 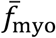 the mean force per myosin II head) (Figure 4G and STAR methods).

The net ring tension *T* sums the tensions of all filaments passing through a given cross-section (Figure 4D). We derived an approximate formula,

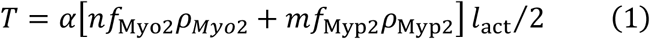

where a Myo2 node or Myp2 cluster contains *n* = *m* = 16 heads, ρ_Myo2_ and ρ_Myp2_ are the cluster densities along the ring, *f*_Myo2_, *f*_Myp2_ are the forces per head pulling actin filaments of mean length *L*_act_, and the factor *α* accounts for whiskering, relative actin-myosin motion and the filament distribution width (see STAR Methods). Using experimental myosin densities and actin filament lengths (Courtemanche et al., 2016; Wu and Pollard, 2005) with no fitting parameters, Eq. 1 reproduced simulated and experimental tension profiles (Figure 4A).

### Membrane-anchored nodes move in both directions around the ring

Due to this tension mechanism, nodes are slowly pulled clockwise or counterclockwise around the ring by myosin II (Figure 4C) as seen in live cells (Laplante et al., 2016). A node carries a mean of ~ 1 formin (Courtemanche et al., 2016; Wu and Pollard, 2005) and hence one filament pulled by myosin. Simulations accurately reproduced the experimental bimodal distribution of lateral node velocities in the membrane, (Figure 4H).

Firm anchoring to the plasma membrane is required for this tension mechanism to work, so the actin filaments become tense when pulled by myosin II (Figure 4C). Indeed, this condition is satisfied: the mean node speed 22 ± 11 nm s^−1^ (*n* = 60 rings, 10 min after constriction onset) is well below the load-free sliding velocity of myosin II ~ 240 nm s^−1^ (Table S1). With artificially lowered node anchor drag coefficient, ring tension decreased ~ 40% for drag extrapolated to zero, in accordance with a simple analytical model (Figure 4I and STAR methods). The residual tension is a second “fixed filament” contribution due to nodes hosting like-oriented filaments forming connected circular chains around the ring.

### Contractile rings have a dense actomyosin core and a dilute actomyosin corona

Cross sections of simulated rings had two zones. At constriction onset the ~ 130 nm thick actin bundle had ~ 47 filaments in its cross-section, in the range of 14-60 filaments reported from cryoET (Swulius et al., 2018). The bundle had a high-density central 100 nm core of ~ 39 filaments, surrounded by a more dilute corona of ~ 5-10 filaments filling a ~ 3-fold greater area (Figure 5A). Myo2 was bound only to core actin filaments and the Myo2 cluster size defined the core diameter, while Myp2 bound all ~ 47 filaments, spanning the entire bundle. Hence the myosins overlap radially, but the median Myp2 location is 26 nm further from the membrane than Myo2, as seen experimentally (Laplante et al., 2015; McDonald et al., 2017), (Figures 5A, B).

**Figure 5.**
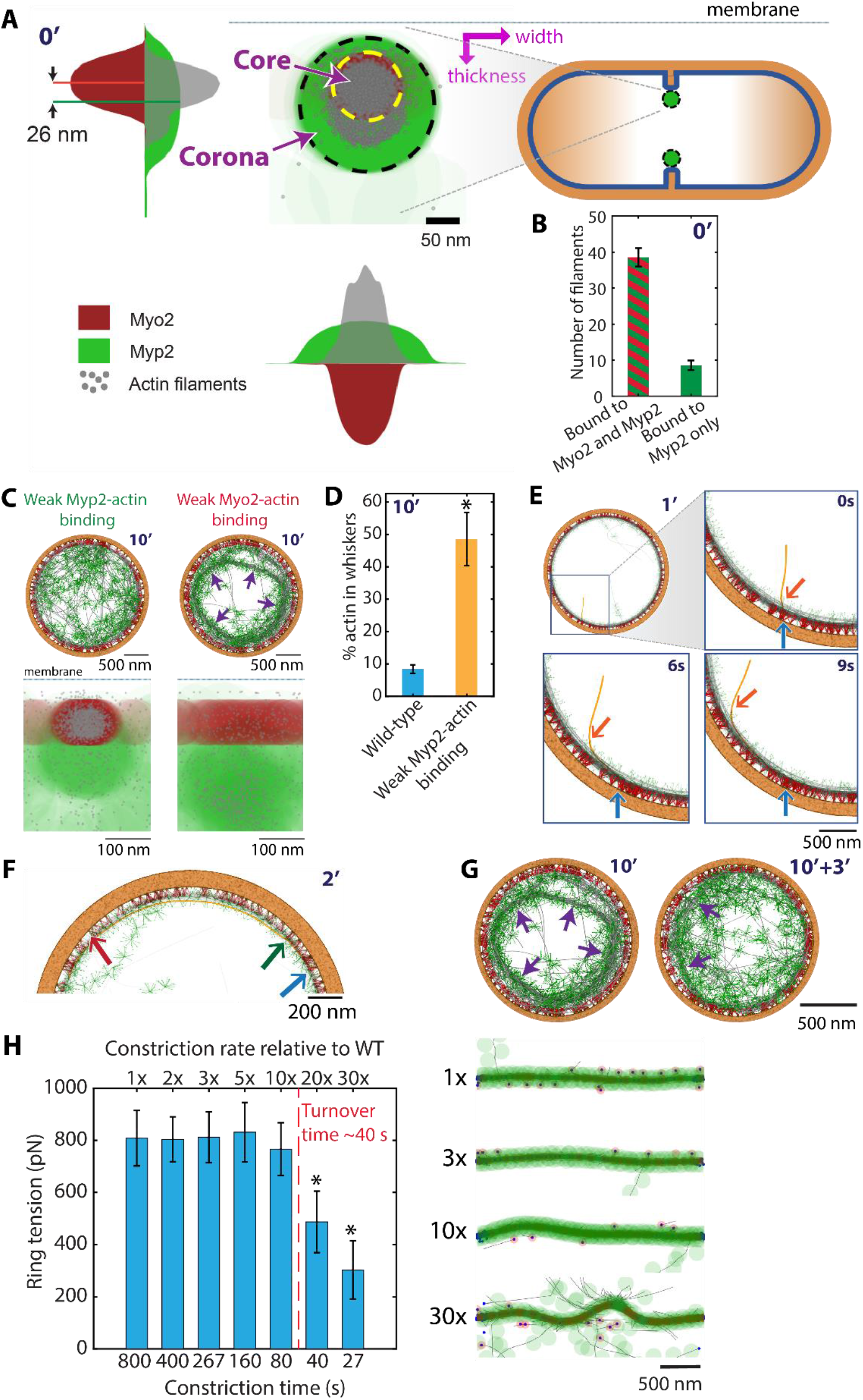
Contractile rings have a dense actomyosin core and a dilute corona. **(A)** Localizations of Myo2 heads, Myp2 heads and actin filaments in the cross-section of a ring at constriction onset. Projection of all cross-sections around the ring are shown. Myo2 and Myp2 organize a dense actin bundle core and a more dilute outer corona, respectively. Indicated core and corona boundaries are circles of radius the Myo2 ellipsoid or Myp2 cluster width, respectively, centered on the median Myo2 or Myp2 head location. Density profiles across the ring width and thickness are plotted (arbitrary units, same units for Myo2 and Myp2). **(B)** Fraction of filaments bound to both Myo2 and Myp2 (core) or Myp2 only (corona). n = 60 rings 10 min after constriction onset, mean ± SD. **(C)** Weakening Myo2- or Myp2-actin binding leads to loss of the core or corona, respectively. The actin-Myp2 or actin-Myo2 filament-cluster unbinding threshold was lowered from 30 pN to 10 pN (left) or from 40 pN to 10 pN (right). Cross-sections are shown in the bottom row. **(E)** In rings with weakened Myp2-actin binding (10 pN threshold) the fraction of actin in whiskers increased ~ 6-fold due to corona loss (n = 10 rings, mean ± SD; * *p* < 0.05). **(F)** Myosin II-mediated zippering (orange arrow) of a newly nucleated actin filament, 1 min after constriction onset. The filament was nucleated by a formin (blue arrow), captured by Myp2, then entrained into the core by Myo2. **(G)** An actin filament (orange) meandering between core and corona, 2 min after constriction onset. Core-corona transition points (green and red arrows) and barbed end (blue arrow) are indicated. **(H)** Ring organizational defects can be long-lived. Using an initial condition with bridges (ring of panel **(C)**), after 3 min the organization is partially but not fully self-repaired (arrows). **(I)** Functional constricting rings require component turnover to be faster than constriction. When constriction rates are artificially increased beyond the turnover rate, ring tension is reduced (n = 10 rings, error bars SD, *p<0.05) and organization drastically disrupted with buckled rings (360^0^ merged in-plane side-view of 1μm radius rings, right). Constriction time defined as −*R*_ring_/(*dR*_ring_/*dt*).

Thus, the core and corona are maintained primarily by Myo2 and Myp2, respectively. Indeed, in simulations with weakened Myo2-actin binding the dense core was abolished: most of the bundle unanchored from the membrane into straight Myp2-decorated bridges (Figures 5C (arrows) and S4). In simulations with weakened Myp2-actin binding the corona was lost, the core became less dense, and ~ 1/2 the ring’s actin was in whiskers (Figures 5C, D and S4).

### Continuous reassembly maintains the organization and tension of the ring

How is this contractile ring organization created and maintained? A long-standing question is how contractile rings remain functional as they constrict, given components are constantly shed (Schroeder, 1972; Wu and Pollard, 2005). Simulations revealed a mechanism of continuous reassembly: as rings constricted, dissociating components were continuously replaced by incoming components that self-assembled into the ring organization without supervision (Stachowiak et al., 2014). Self-assembly processes included myosin II-mediated zippering of formin-nucleated actin filaments into the bundle, entrainment of incoming nodes, meandering of growing filaments that welded the core and corona, and actin-dependent recruitment of Myp2 (Figures 3D & 5E, F).

The continuous renewal of organization enabled rings to constrict without losing functionality. This mechanism required that the renewal time (~ 1 min) be much less than the constriction time (~ 25 mins), so ring components experienced almost no change in ring length before being replaced. Then the ring could rebuild itself at each progressively shorter length without disruption. Indeed, the organization was violently disrupted in simulations with constriction rates artificially boosted to exceed turnover rates. Nodes aggregated, rings buckled and the tension was dramatically lowered or negative (Figure 5H).

The actomyosin bundle was self-sustaining: once initiated, it provided a platform for further zippering and for Myp2 to bind and participate in zippering-anchoring (Figure 3D). This history dependence enabled the ring to maintain a stable tension-generating organization, but also meant that radical organizational alterations were long lived. Artificially imposed partially unanchored bridges remained kinetically locked-in by Myp2-mediated zippering, without assistance from Myo2 (Figure 5G).

### Simulated *myo2-E1* rings reproduce experimental tension loss and bridging defects

Next we simulated the temperature-sensitive *myo2-E1* mutant, which has ~ 2/3 reduced ring tensions (Figure 2) and severely disrupted organization with sections of the ring detaching from the membrane into straight bridges containing Myp2 (Laplante et al., 2015).

Myo2-E1 has minimal ATPase activity, binds actin weakly, and fails to translocate actin (Lord and Pollard, 2004; Stark et al., 2013). Accordingly, we simulated Myo2p-E1 with zero pulling force on actin, and a lower unbinding threshold (12 pN versus 40 pN WT) which reproduced the experimental bridging phenotype.

Simulations reproduced both features seen in live cells (Figure 6A). The mean tension 190±50 pN was similar to the experimental mean of 220±200 pN (Figure 6B) and tensions were ~ 75 % lower than simulated WT tensions, similar to the ~ 60% reduction seen experimentally (Figure 6C). Due to the weakened Myo2-actin binding, centripetal Laplace forces pulled tense actin filaments away from Myo2 clusters, so entire ring sections formed straight actin bridges containing Myp2 only, anchored by a few tenuous attachments at bridge ends mediated by barbed-end-anchored actin filaments (Figure 6A). Consequently, 10 mins into constriction the actin filament core was severely depleted, with ~ 28% of actin in bridges or whiskers, 3-fold more than in WT (Figure 6D).

**Figure 6.**
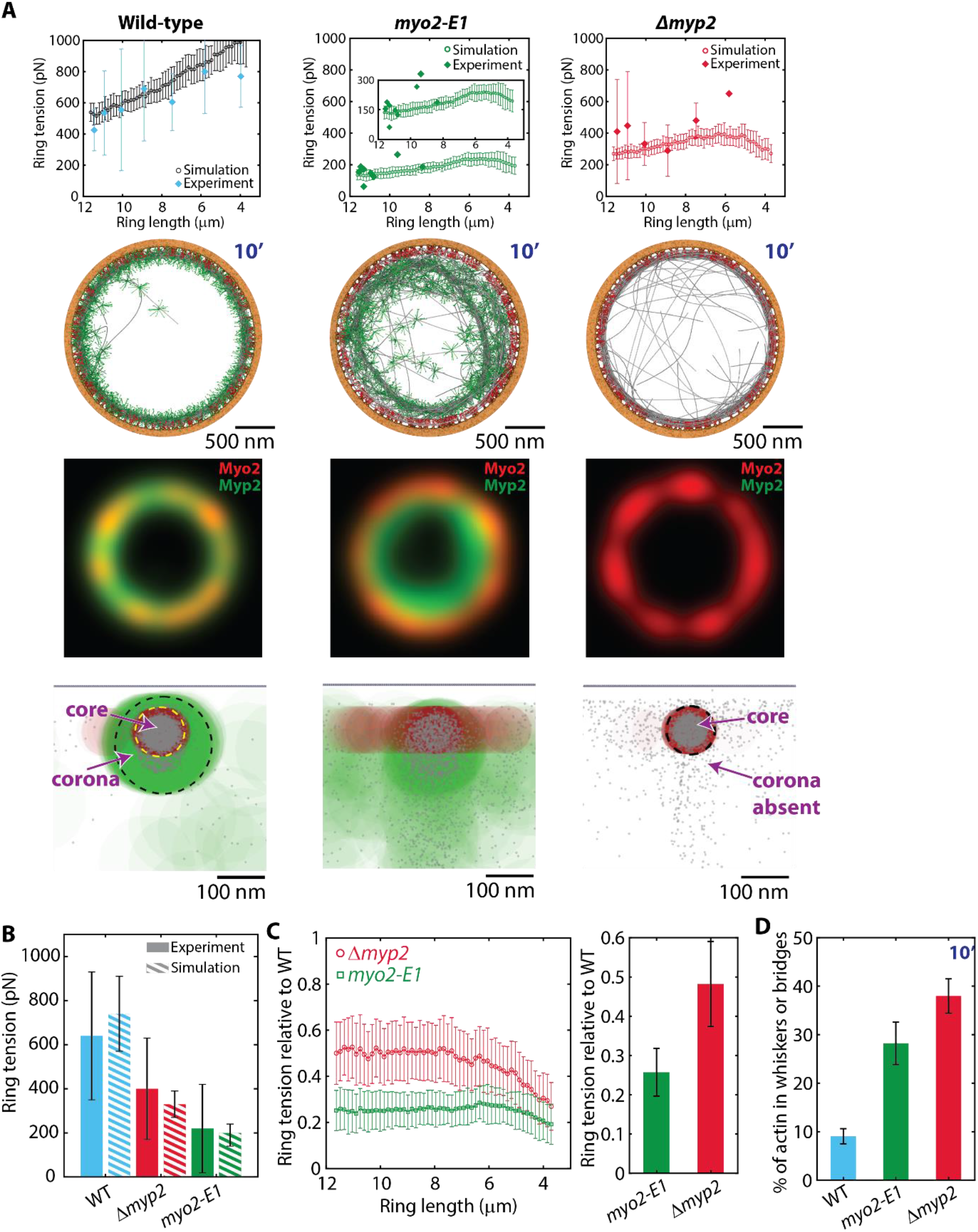
Simulations reproduce tension loss and organizational defects in contractile rings of *myo2-E1* and *Δmyp2* mutants. **(A)** Ring tensions in simulations of contractile rings in WT, *myo2-E1*, and *Δmyp2* cells (n = 60 per simulation condition, mean ± SD) compared to the experimental tensions of Figure 2C. Simulation snapshots show bridging instabilities in *myo2-E1* rings and loss of corona in *Δmyp2* rings (second row). Simulated fluorescence microscopy images (2D Gaussian point spread function, width 310 nm, third row) and cross-section component localizations (fourth row) of the rings shown in the second row. **(B)** Mean simulated and experimental tensions of **(A)**. Error bars are SD. **(C)** Ring tensions of simulated *myo2-E1* and *Δmyp2* rings of **(A)** relative to wild type simulated ring tensions of **(A)**, versus ring length. Mean relative values over all lengths (right). **(D)** Mean fraction of actin in whiskers or bridges in the simulated rings of (A) 10 min after constriction onset.

### Simulated *Δmyp2* rings lack the corona and have reduced tension

We simulated rings in *Δmyp2* cells by omitting Myp2 (Figure 6A). At constriction onset tensions were ~ 50% the WT value, decreasing to ~30% by the end of constriction (Figure 6C), with a mean of 330±60 pN, Figure 6B. By comparison, we measured a mean *Δmyp2* tension in live cells of

~400±200 pN, 65% of the WT value, Figure 6B.

In simulated *Δmyp2* rings the actin bundle had only a reduced density core of ~ 27 filaments (Figure 6A) and the corona was absent, as expected since Myp2 served as its scaffold in WT rings (Figure 5A,B). About 1/3 of actin filaments were chaotically oriented whiskers (Figure 6D).

The reduced ring tension was due in part to the absence of Myp2 pulling forces, and in part to the large whisker population which lowered the mean length of functional actin filaments belonging to the bundle. For the same amount of myosin II, shorter filaments produce lower tension (Eq. (1)).

## Discussion

The contractile actomyosin ring generates tension for cell division, yet this most fundamental property has rarely been measured. In echinoderm embryos, Rappaport (Rappaport, 1967, 1977) and Hiramoto (Hiramoto, 1975) used deformation of microneedles and ferrofluid droplets, respectively, to measure inward radial furrow forces of ~10–50 nN. However these were net forces, including opposing cortical forces. Yoneda and Dan estimated ring tensions of similar magnitude in dividing echinoderm eggs using an approximate force balance at the furrow (Yoneda and Dan, 1972) but the method assumed uniform cortical tension and a symmetrical cell shape with spherical caps, inconsistent with experiment (Hiramoto, 1967, 1975).

These historic studies were complicated by interference from contiguous actomyosin cortex. Here we isolated contractile rings in fission yeast protoplasts free of cortex or cell wall, so ring tension was balanced entirely by passive plasma membrane tension (Figure 1). We measured ring tensions ~ 600 pN, increasing ~ 2-fold as rings constricted, to which Myo2 and Myp2 contributed ~60% and ~30-40 %, respectively (Figure 2). Acting over a cross sectional area ~0.01 μm^2^ (Laplante et al., 2016; Swulius et al., 2018), these tensions produce stresses ~50 nN μm^−2^, similar to the estimated **>**8 nN μm^−2^ exerted by echinoderm egg rings (Rappaport, 1977). Thus, animal cell contractile rings, though much larger, may use similar tension-producing mechanisms.

### Constriction rate is not a proxy for ring tension

In the absence of ring tension measurements, normal constriction rate is routinely taken to signify a normal contractile ring (Laplante et al., 2015; Piekny and Mains, 2002). We find this inference is unreliable. Rings in *myo2-E1* cells constrict at the normal rate (Laplante et al., 2015) in spite of severe tension loss (Figure 2), while both tension and constriction rate (Laplante et al., 2015) are reduced in *Δmyp2* cells. Further, ring tension *T* increased as rings constricted (Figure 2B), so the inward force ~*T*/*R*_ring_ increases dramatically with decreasing radius *R*_ring_, yet constriction rates show no increase as constriction progresses (Thiyagarajan et al., 2015).

Thus, the constriction rate has no simple relation to ring tension. This is not surprising, given the tensions we measured are far too weak to oppose turgor pressure or to mechanically perturb the septum that grows inwards in the wake of the constricting ring (Thiyagarajan et al., 2015). The ring constriction rate is thus limited by the rate of growth of the septum to which it is indirectly attached. As septation is mediated by glucan synthases at the furrow (Cortes et al., 2007), our results suggest Myo2 affects their mean activity but Myp2 does not. If the glucan synthases are mechanosensitive, ring tension may regulate local septum growth rates to ensure proper septum closure (Pollard and O’Shaughnessy, 2019; Thiyagarajan et al., 2015).

### A sliding node mechanism generates contractile ring tension

How does the irregular node-based organization of the contractile ring lead to tension? The critical feature is that nodes host actin filaments, whose barbed ends are thus firmly anchored to the membrane thanks to the small lateral node mobility in the membrane (Vavylonis et al., 2008). Thus when myosin II pulls actin filaments, host nodes slide slowly clockwise or counterclockwise around the ring and the filaments are made tense (Figure 7A). The myosin doing the pulling belongs to other nodes. Thus, in the disordered steady state ring, oppositely oriented nodes within range are continuously drawn together without ring shortening, which occurs on a much longer ~ 25 min timescale. This is in contrast to sarcomere shortening, which shortens muscle.

**Figure 7.**
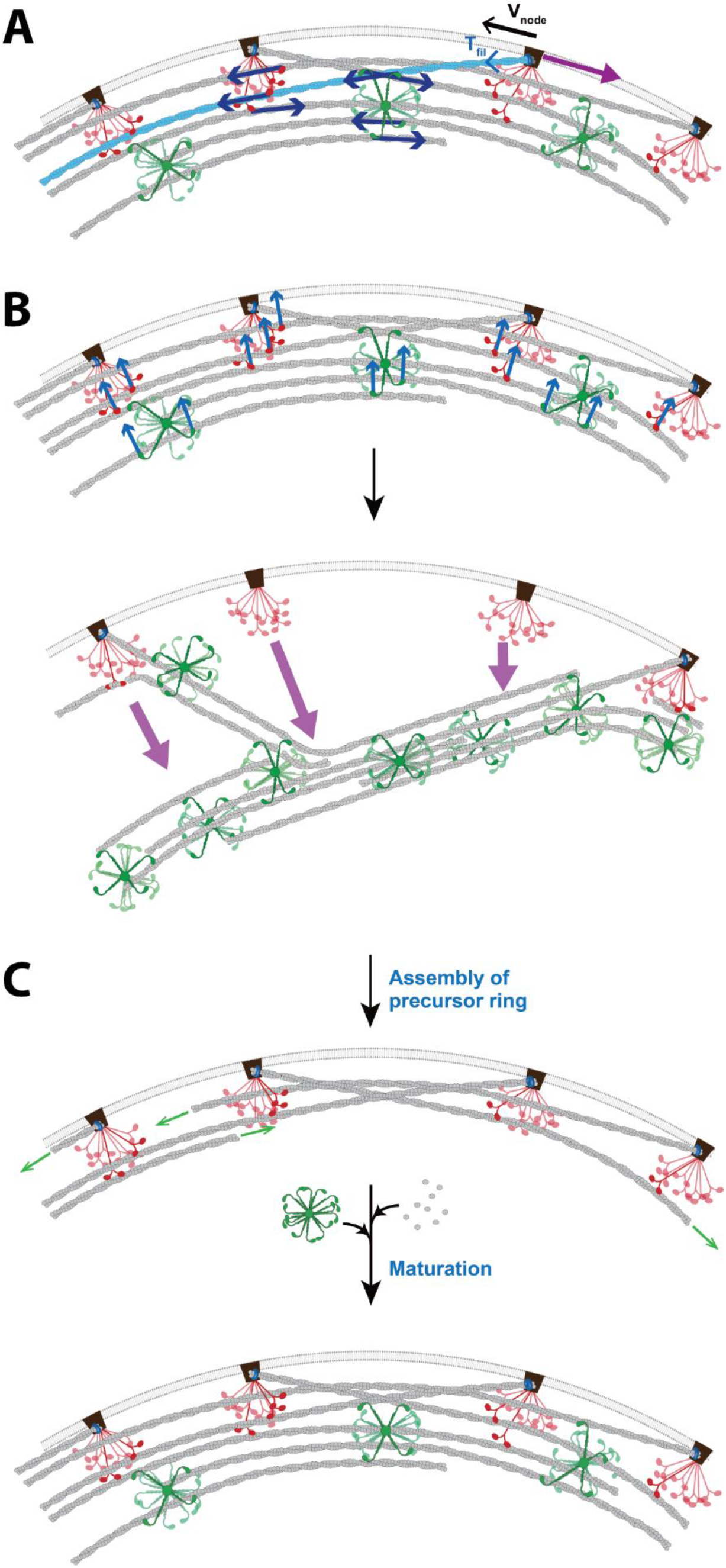
Model of the fission yeast contractile ring: assembly, anchoring and tension production. **(A)** Sliding node tension generation mechanism. Myo2 (red) and Myp2 (green) pull actin filaments whose barbed ends are anchored to the membrane by nodes. Large node-membrane drag forces (purple arrow) ensure nodes slide slowly (*ν*_node_) and filaments develop tension (*T*_fil_). Unanchored Myp2 contributes due to the balanced bundle polarity. **(B)** Myo2 and Myp2 cooperatively anchor and crosslink the actin filament bundle (forces, blue arrows) protecting the ring from bridging instabilities driven by the energetic preference of tense filaments to straighten. In *myo2-E1* cells Myo2-actin binding is too weak to prevent bridging. **(C)** Myo2 assembles the precursor ring from nodes. During maturation to the constriction-ready ring, actin filaments elongate and recruitment of unanchored Myp2 allows the ring to thicken.

Clockwise and counterclockwise node motions observed in live cells (Laplante et al., 2016) have a velocity distribution close to that predicted by simulations (Figure 4H), with a mean speed well below the myosin II gliding velocity, consistent with substantial filament tensions (Table S1). A similar mechanism is used during ring assembly, when actin filaments are made tense as node pairs are pulled together from a broad band (Vavylonis et al., 2008). Constriction of partially anchored rings in fission yeast cell ghosts is quantitatively consistent with tension being generated by barbed-end anchoring of actin filaments to the membrane (Mishra et al., 2013; Wang and O’Shaughnessy, 2019).

The trends in contractile ring scaleup in larger animal cells suggest the broad mechanism based on barbed-end-anchored actin may be conserved. Ring length and width increase with cell size, but ring thickness remains several hundred nm, similar to fission yeast (O’Shaughnessy and Thiyagarajan, 2018). Thus, in larger cells the plasma membrane remains accessible for barbed end anchoring of actin filaments lying parallel to the plasma membrane.

### Myo2 and Myp2 make force, create ring organization and protect against instability

In simulations Myo2 and Myp2 had broad, complimentary roles in the ring. Beyond force production, they helped create and protect the ring organization. The anchored nodes radially positioned Myo2 heads rather precisely, so actin and Myo2 cooperatively condensed into a dense core of ~ 30-35 actin filaments with well-regulated spacing from the membrane (Figure 5A). This stable Myo2-bundled core allowed the unanchored Myp2 to roam beyond the range of Myo2 and entrain ~ 5-10 more filaments in a loosely packed outer corona (Figures 5A & 7B).

Myo2 also serves to anchor the actin bundle to the membrane, protecting the ring from bridging instabilities (Figure 7B). These are by far the biggest threat to contractile ring integrity, originating in the enormous energetic preference for a high tension curved bundle to shorten by straightening. In *myo2-E1* cells with weakened actin binding, rings featured straight unanchored Myp2-containing bridges (Laplante et al., 2015). *myo2-E1* simulations yielded a remarkably similar phenotype, with sections unanchoring into straight bridges (Figure 6A).

Why is Myo2 the ring’s principal anchor? Depending on the duty ratio, up to ~ 250 actin-bound Myo2 heads μm^−1^ of ring (Wu and Pollard, 2005) oppose the Laplace forces *T*/*R* ~ 220 pN μm^−1^. Duty ratios of NMIIA and NMIIB in animal cells are high under load (Kovacs et al., 2007), so Myo2 may exhibit similar mechanosensitivity. We estimate each Myo2 head must support ~ 4 pN (see STAR methods), within the ~ 6-10 pN unbinding threshold measured for muscle myosin II (Guo and Guilford, 2006; Nishizaka et al., 1995).

Furthermore, Myo2 is uniquely qualified as an anchor in the *S. pombe* ring. As a membrane-anchored motor protein using actin filaments for tracks, it can accommodate the constant elongation, fragmentation and sliding of filaments without unbinding. Passive actin-binding anchors would rapidly unbind from such dynamic filaments. Indeed, *α*-actinin ain1 contributed insignificantly to bundling or ring tension (Figure S7) consistent with *ain1* deletion cells assembling and constricting rings normally (Wu et al., 2001) (see STAR methods).

More generally, the processivity of myosin may lend itself to unexpected anchoring roles. Anchoring of actin to the plasma membrane is thought accomplished by myosin VI in mammalian sensory hair cells (Altman et al., 2004; Self et al., 1999) and by myosin I during clathrin-mediated endocytosis in budding yeast (Pedersen and Drubin, 2019).

### Myo2 and Myp2 have complimentary roles in assembly of the constriction-ready ring

To assemble the constriction-ready fission yeast ring, a precursor ring is first assembled from Myo2-containing nodes, followed by a ~25 min maturation phase when Myp2 is recruited (Wu and Pollard, 2005) (Figure 7C). We assumed Myp2 is not directly anchored to the membrane, suggested by several findings. Localization of Myp2 to the ring depends on actin filaments (Laplante et al., 2015; Takaine et al., 2015) and Myp2 lies >100 nm from the plasma membrane (McDonald et al., 2017), further than Myo2 (Laplante et al., 2015). Myp2 is present in unanchored straight bridges in mutants of Myo2 (Laplante et al., 2015) and in mutants of ADF/cofilin when Myo2 was shown to be left behind at the plasma membrane (Cheffings et al., 2019).

Why then is an unanchored myosin II recruited during maturation? Presumably, a major goal of maturation is to boost ring tension. This can be achieved by adding myosin or by elongating the actin filaments already present (Eq. (1)). Indeed, both measures are taken during maturation, when ~2000 Myp2 molecules augment the ~3000 Myo2 molecules in the precursor ring, and the polymerized actin increases ~40% with no change in the number of Cdc12 formins (Courtemanche et al., 2016; Wu and Pollard, 2005). Using the best fit stall forces (Table S1) in eq. (1) gives *T*~(*ρ*_myo2_ + 0.6 *ρ*_myp2_) *L*_act_, so these maturation measures increase ring tension ~2-fold.

However, elongating actin filaments in this fashion thickens the ring beyond the reach of Myo2, a significant technical challenge (Figure 7C). Unanchored Myp2 provides an ideal solution, able to roam from the membrane and bundle the additional actin into a corona while contributing to tension despite being unanchored.

Thus, the myosin II isoforms have complimentary roles during assembly of the ring into its constriction-ready form, just as they do during constriction. Membrane-anchored Myo2 is ideal for assembly of the nodes into a precursor ring, whereas unanchored myosin II would be useless for this task as nodes link up using just one or two filaments (Vavylonis et al., 2008). Once assembled, unanchored Myp2 is ideal to mediate thickening of the ring and a ~2-fold tension increase during maturation, a significant “beefing up.”

## Supporting information

STAR Methods

## Acknowledgements

This work was supported by National Institute of General Medical Sciences of the National Institutes of Health under award number R01GM086731 to B.O’S. and R01GM026132 to T.D.P. The content is solely the responsibility of the authors and does not necessarily represent the official views of the National Institutes of Health. H.F.C. is a Merck LSRF fellow. We thank C. Laplante and R. Arasada for yeast strains (all Pollard lab stock) and helpful discussions, J. Nikolaus for assistance with cell culture and the micromanipulator setup, and Dong An for assistance with figures.

## AUTHOR CONTRIBUTIONS

B.O’S. conceived the study, B.O’S., T.D.P., E.K. and H.C. designed the experiments, B.O’S.,S.W., S.T., and Z.M. designed the model, H.C. performed the experiments, S.W., Z.M., and S.T. performed the simulations, B.O’S., H.C., S.W., Z.M., and S.T. analyzed the data. B.O’S, T.D.P., S.W., H.C, S.T., and Z.M. wrote the manuscript.

