## Supplementary material for "Myosins generate contractile force and maintain organization in the cytokinetic contractile ring": STAR Methods

### CONTACT FOR REAGENT AND RESOURCE SHARING

### EXPERIMENTAL MODEL AND SUBJECT DETAILS

#### Yeast Strains

Fission yeast *S. pombe* strains used in this study are listed in KEY RESOURCES TABLE. Cells were grown in an exponential phase at 25°C in YE5S rich liquid medium.

### METHOD DETAILS

#### Preparation of protoplasts

Removal of fission yeast cell wall was performed as described (Stachowiak et al., 2014) with minor modifications. Protoplasts were generated by shaking for 30-60 min at 25°C in EMM5S supplemented with 0.8 M sorbitol containing protoplasting enzyme cocktail: 5 mg/mL Lysing Enzymes (Sigma-Aldrich) and 5 mg/mL Yeast Lytic Enzymes (MP Biomedicals). Such enzymatic digestion of the predominantly β-glucan cell wall generated spherical protoplasts of diameter ~ 5-7 μm (Jochova et al., 1991; Kopecka, 1975; Mishra et al., 2012; Stachowiak et al., 2014). Protoplasts were collected by centrifugation at 2000g for 30 sec and washed 3x in PBS supplemented with 0.8 M sorbitol. The protoplast surface was labeled by biotinylation with 5 mg/mL Biotin-PEG-NHS, 3kDa (Nanocs) in PBS supplemented with 0.8 M sorbitol, followed by incubation with 5μM fluorescent Alexa555- or Alexa647-streptavidin (Life Technologies) for 10 min at 4°C. Cells were resuspended in imaging media (EMM5S + 0.8 M sorbitol) containing 2x diluted protoplasting enzyme cocktail. The osmolarity of solutions was measured with a Micro-Osmette osmometer (Precision Systems Inc.) and adjusted with D-sorbitol (Fischer Scientific) to equalize osmolarity of solutions. The protoplasts regrew little cell wall up to one hour after protoplasting (Figure 1A).

### Micropipette aspiration

The membrane tensions of protoplasts were measured using micropipette aspiration (Hochmuth, 2000; Stachowiak et al., 2014) in ES buffer (EMM5S, 0.8 M sorbitol, 0.2% BSA (Sigma), 2.5 mg/mL Lytic enzymes, 2.5 mg/mL Lysing enzymes) within 1 h of completion of the enzyme treatment. Suction pressure in the aspirating micropipette was controlled using a hydrostatic system. The membrane tension $\sigma_1$ in the aspirated lobe was determined from Laplace's law, $\sigma_1 = \Delta P_{\mathrm{pip}}/2\left(R_{\mathrm{p}}R_1/\left(R_1 - R_{\mathrm{p}}\right)\right)$ (Hochmuth, 2000). Ring tension $T$ was determined by measurement of the furrow angles $\theta_1, \theta_2$ and the membrane tension $\sigma_1$ using $T/R_{\mathrm{ring}} = \sigma_1 \cos\theta_1 + \sigma_2 \cos\theta_2$. Here $\sigma_2 = \sigma_1 R_2/R_1$ is the membrane tension in the other lobe. See section "Measurement of ring tension" for details.

### Microscopy and image analysis

Imaging was performed using a spinning disc confocal system (Volocity, PerkinElmer, Waltham, MA) on an inverted microscope (Eclipse TE2000-E, Nikon, Melville, NY) with a Hamamatsu Orca-ER camera and 100x/1.45 Plan Apo oil-immersion objective (Nikon), using proper excitation/emission filters and solid state laser lines for given fluorescent probes. Image analysis was performed using ImageJ (http://rsb.info.nih.gov/ij/).

### Measurement of ring tension

In Stachowiak et al. (Stachowiak et al., 2014) we estimated the ring tension in fission yeast protoplasts from a force balance at the furrow, using a value for the membrane tension obtained by averaging measurements on cells without rings. We assumed that in the absence of a cell wall the surface stresses due to furrowing by the cytokinetic ring are supported by the plasma membrane alone, since fission yeast lacks an extended actin cortex (Kovar et al., 2011; Marks and Hyams, 1985). Here we advanced the method by measuring the membrane tension in individual cells with a ring, and using this value in the force balance for that cell.

Briefly, we aspirated one lobe of the furrowed protoplast and obtained the membrane tension from the applied suction pressure and the geometry of the cell using Laplace's law. The ring tension was then inferred using a force balance at the furrow that depends on the membrane tension in the two lobes, and the geometry of the furrow and the lobes. This is an advance on the method used in our previous study (Stachowiak et al., 2014) where an interphase membrane tension was measured, not on the same cell as that to which a force balance was applied. The previous measurements likely underestimated the ring tension as the measured membrane tension of unfurrowed protoplasts is significantly smaller than the membrane tension of furrowed protoplasts (Figure S1A). Our tension measurements were consistent for three cell lines expressing ring markers Rlc1p-tdTomato, Rlc1p-3GFP or CHD-GFP.

Force balance at the furrow gives ring tension. The fission yeast protoplast roughly behaves as a fluid with an interfacial tension with the external environment corresponding to the cell's membrane tension. This is exemplified by the uniform curvature of the cell surface following removal of the cell wall, resulting in a spherical shape in the absence of a cytokinetic ring or two semispherical lobes intersecting at a cytokinetic ring in furrowed protoplasts (Figure 1). The ring is significantly thinner than its radius, so the forces at the furrow generated by the ring can be examined within the plane of the ring (Figure 1B).

The force balance in the plane of the ring is described by two opposing forces: 1) an inward force per unit length along the ring due to ring tension ($T/R_{\text{ring}}$), and 2) a resistive force generated by the membrane tension along the outward radial direction (Figure 1B). We can thus determine ring tension (i.e., the tension within the actomyosin bundle due to myosin contractility) from this force balance

$$T/R_{\text{ring}} = \sigma_1 \cos\theta_1 + \sigma_2 \cos\theta_2,$$

where $R_{\text{ring}}$ is the radius of the ring, $\sigma_1$ and $\sigma_2$ are the membrane tensions of the two lobes, and $\theta_1$ and $\theta_2$ are the angles made by the lobes at the furrow (Figure 1B). Yoneda and Dan presented a similar force balance for dividing echinoderm eggs, but with a symmetry axis in the ring plane ($\theta_1 = \theta_2$) (Yoneda and Dan, 1972). The angles made by the lobes at the furrow are difficult to measure accurately in echinoderm embryos and other model organisms, as they do not strictly adopt spherical lobes, presumably due to gradients in cortical tension from the cell poles to the furrow. In contrast, furrowed protoplasts very closely approximate two spherical lobes, such that the angles can be measured accurately by measuring the radii ($R_1$, $R_2$) and the ring radius $R_{\text{ring}}$.

In this force balance, we neglect the cytoplasmic viscous forces, because a simple calculation shows that they are orders of magnitude smaller than the inward force per unit length due to ring tension, as follows. We approximate the ring as a cylinder of diameter 2w = 125 nm and max length $L_{Ring}$ = 20 μm. The drag coefficient per unit length of this cylinder is approximately (Broersma, 1960) $\zeta \sim 4\pi\nu_{\text{cyt}} / [\ln(2L_{\text{Ring}}/w) - 1.13] \sim 1$ pN $\mu\text{m}^{-2}$ s, where $\nu_{\text{cyt}} \sim 0.3$ Pa · s is the cytoplasmic viscosity (Alexander and Rieder, 1991). Therefore, with a constriction rate of ~ 7 nm $s^{-1}$ (~ 20 μm initial length, Figure S1D, constriction time ~ 50 min, (Stachowiak et al., 2014)), protoplast rings are subject to a cytoplasmic drag force ~ 7 fN per micron along the ring, 4 orders of magnitudes smaller than the inward force per unit length due to ring tension, $T/R_{\text{Ring}} \sim 100$ pN $\mu\text{m}^{-1}$.

The five observables ($R_{\text{ring}}$, $\theta_1$, $\theta_2$, $\sigma_1$, and $\sigma_2$) required to calculate ring tension $T$ from the force balance equation were extracted from a combination of imaging and micropipette aspiration as follows:

1) *Ring radius, $R_{\text{ring}}$*. To measure $R_{\text{ring}}$, the z-plane passing through the center of the ring and with its normal perpendicular to the long axis of the cell was used as the confocal imaging plane. In this plane, fluorescence from Rlc1p-tdTomato or Rlc1p-3GFP appears as two spots. The distance between these spots is the diameter of the ring, $2R_{\text{ring}}$. The ring radius was confirmed by generating a single face-on view image of the ring using the ImageJ Reslice function, followed by fitting the myosin fluorescence to a circle.
2) *Contact angles at furrow ($\theta_1$, $\theta_2$)*. The furrow angles are determined from the relations, $\sin\theta_1 = R_{\text{ring}}/R_1$ and $\sin\theta_2 = R_{\text{ring}}/R_2$. The angles are obtained as $\theta_{\text{i}} = \sin^{-1} R_{\text{ring}}/R_{\text{i}}$ or $\theta_{\text{i}} = \pi - \sin^{-1} R_{\text{ring}}/R_{\text{i}}$ for $i = 1, 2$ depending on the geometry of the furrow. We measured $R_1$ and $R_2$ by fitting the membrane fluorescence marker (Alexa555-streptavidin or Alexa647-streptavidin) to circles in the confocal z-plane where the radii are maximal,

corresponding to the plane passing through the center of each lobe. In some cases where the cell's long axis is not parallel to the imaging plane, these are different confocal slices for the two lobes.

3) *Membrane tension of aspirated lobe,* $\sigma_1$. Membrane tension $\sigma_1$ in one lobe of the furrowed protoplast with radius $R_1$ was directly measured by micropipette aspiration, by measuring the suction pressure $\Delta P_{pip}$ needed to aspirate a membrane tongue of length equal to the pipette radius $R_p$. From Laplace's law, $\sigma_1 = \Delta P_{pip}/2(R_p R_1/(R_1 - R_p))$[3,4].

4) *Membrane tension of non-aspirated lobe,* $\sigma_2$. Laplace's law for a spherical geometry $\Delta P = 2\sigma/R$ shows that the membrane tension in each lobe scales by the lobe radius $\sigma_1/R_1 = \sigma_2/R_2$, assuming uniform internal pressure throughout the protoplast. We found that the membrane tension in each lobe varies with local radius (Figs. 1D-F), consistent with a uniform internal pressure. We were able then to calculate $\sigma_2$ from the measurements of $R_1$ and $R_2$ (from membrane fluorescence images), and of $\sigma_1$ (from micropipette aspiration).

The simple geometry of the furrowed protoplast (Figure 1B) thereby readily allowed us to measure ring tension $T$ from measurements of $R_1$, $R_2$, $R_{ring}$, and $\sigma_1$ in static confocal images of aspirated protoplasts using the force balance equation. Our measurement relies upon the assumption that membrane tension is the only significant resistance to ring tension in protoplasts. Our calculations suggest that cytoplasmic viscous resistance, which is a possible source of error, is likely to be small in comparison to the ring tensions measured in protoplasts.

Ring tension measurement over time. To measure ring tension $T$ as a function of time (Figure 3), we first obtained the aspiration pressure $\Delta P_{pip}$ required to aspirate a tongue of length $R_p$ for one lobe of the furrowed protoplast. We maintained the aspiration pressure $\Delta P_{pip}$ while imaging the protoplast at 2-5 minute intervals. As the ring constricts, it slides towards the smaller lobe, causing the larger, aspirated lobe radius to increase. While this would be expected to change the required aspiration pressure to maintain the tongue length on long time scales, the change in lobe radius was small enough that we did not observe an appreciable change in tongue length during the observation time window of 20-40 minutes.

**Simulation of the contractile ring**

We built a molecularly explicit simulation of the fission yeast contractile ring, incorporating experimental measurements as much as possible. The simulation was implemented in MATLAB (MathWorks), and determines the positions, velocities, and amounts of ring constituents as the ring constricts (see section "Running the simulation" for details). The amounts and organizations of the ring components have already been described in the main text. Here we describe in detail the interaction rules between components.

Amounts of ring components. (i) Myosin II. 2900 Myo2 and 2000 Myp2 molecules are present at the onset of constriction (Figure 3B and 3D of (Wu and Pollard, 2005)). We obtained the densities $\rho_{\mathrm{Myo2}}$ and $\rho_{\mathrm{Myp2}}$ at any given time during constriction by fitting data points obtained from (Wu and Pollard, 2005) with interpolating splines. (ii) Formin Cdc12. 200 dimeric molecules are present. at the onset of constriction, decreasing slightly until a late, sharp decrease (Figure 5D of (Courtemanche et al., 2016)). We obtained the density $\rho_{\mathrm{Cdc12}}$ at any given time during constriction by fitting data points obtained from (Courtemanche et al., 2016) with an interpolating spline. (iii) Actin filaments. The number of actin filaments in the simulation is set by the number of formin Cdc12p dimers; the time-dependent mean length is calculated as the total filament length from (Courtemanche et al., 2016) divided by the number of filaments. (iv) Actin crosslinker α-actinin. 250 α-actinin dimers are present at the onset of constriction (Wu and Pollard, 2005), and we assume constant α-actinin density during constriction.

Organization of ring components. (i) Heads of constriction nodes, Rng2p and Cdc15p. We modeled these two proteins as a sphere with 70 nm diameter, the spread of Cdc15p in FPALM (Laplante et al., 2016). The centers of these spheres are 40 nm from the membrane, following the center of Rng2 in FPALM (Laplante et al., 2016) (Figure 4A). (ii) Formin Cdc12p are located within the heads of constriction nodes. We assume that formin dimers are centered at the same point as the node heads for convenience, 40 nm from the membrane (Laplante et al., 2016). Because they are within the heads of constriction nodes, we assume they have no excluded volume for simplicity (Figure 4A). (iii) Myo2 dimers. Each node contains 8 Myo2 dimers. Myo2 tails colocalize with the heads of the nodes (Laplante et al., 2016), and hence are not explicitly represented for simplicity. All 16 Myo2 heads are generically represented as a 132 x 102 X 102 nm ellipsoid centered 94 nm from the membrane, centered on a line perpendicular to the plasma membrane and passing through the center of the Rng2p/Cdc15p, with the long axis of the ellipsoid aligned circumferentially along the ring (Figure 4A). Myo2 binds and pulls actin filaments that overlap with this ellipsoid. (iv) Myp2 clusters. As both the heads and tails of Myp2 are >100 nm from the plasma membrane (McDonald et al., 2017), we model Myp2 clusters as unanchored spheres of radius 100 nm, comparable to the unfolded Myp2 tail length ~130 nm (Bezanilla and Pollard, 2000). This choice resulted in a 200 nm thick distribution of Myp2 in simulated rings, similar to the ~175 nm thickness of the Myp2 distribution measured by FPALM (McDonald et al., 2017). Myp2 binds and pulls actin filaments that overlap with this sphere. (v) Actin filaments. Each actin filament is represented as a chain of 100 nm rods. Torsional springs at the hinges of the chain imposes a bending stiffness that corresponds to the experimentally measured persistence length ~ 10 μm (Ott et al., 1993).

Excluded volume interactions. There are excluded volume interactions between the following: (i) Myp2 clusters, (ii) Myo2 clusters, (iii) actin subunits, (iv) Myp2 clusters and node anchors. For the Myp2-Myo2, anchor-Myp2, and Myo2-Myo2 interactions, the excluded volume force is of the form

$$\vec{f} = k_{\text{ex}}(r - r_0)\hat{r} \quad r < r_0,$$

where $r$ is the distance between the particles, $r_0$ is the cutoff distance for the interaction, $\hat{r}$ is the unit vector pointing from one particle to the other, and $k_{\text{ex}}$ is the spring constant of the force. For the actin-actin interaction, the force is given by

$$\vec{f} = k_{\text{act}}\, e^{-r^2/r_0^2}\, \hat{r} \quad r < 2r_0,$$

where $k_{\text{act}}$ is the maximum force, and $r_0$ is the decay length for the force.

Binding of myosin to actin. As stated in the section *Organization of ring components* above, Myo2 molecules are present in membrane-anchored nodes and Myp2 molecules are represented as unanchored clusters. All Myo2 molecules of a node and all Myp2 molecules in a cluster are represented by one ellipsoid or one sphere respectively. Any actin filament that intersects these objects experiences an attractive binding potential (Figure 4B). Thus, each geometric object represents the capture zone of that particular node or Myp2 cluster respectively.

An actin filament segment that intersects a Myo2 ellipsoid experiences an attractive force that points towards the center of the ellipsoid. If more than one segment on a filament are within the capture zone, the capture force is exerted only on the segment closest to the pointed end. The formula for the Myo2-actin binding force is as follows:

$$\vec{f}_{i,\alpha}^{\mathrm{cap}} = \begin{cases} -k_{\mathrm{Myo2}}\{(\vec{d} - d_{Myo2}\hat{d}) - [(\vec{d} - d_{Myo2}\hat{d}) \cdot \hat{\mathrm{T}}_{\mathrm{i}}]\hat{\mathrm{T}}_{\mathrm{i}}\}, & |\vec{d}| \geq d_{Myo2} \\ 0, & |\vec{d}| < d_{Myo2} \end{cases}$$

where $\vec{d} = \vec{r}_i - \vec{r}_\alpha$ is the displacement between the center of the capture zone of Myo2 α and actin subunit *i* with $\hat{d}$ representing the corresponding unit vector, $\hat{T}_i = (\vec{r}_{i+1} - \vec{r}_i)/|\vec{r}_{i+1} - \vec{r}_i|$ is the unit tangent vector of the actin filament at subunit *i*, unless the subunit *i* is at the pointed end, where $\hat{T}_i = (\vec{r}_i - \vec{r}_{i-1})/|\vec{r}_i - \vec{r}_{i-1}|$, and $d_{Myo2}$ equals 25.5 nm, one-fourth of the length of the short axis of the Myo2 ellipsoid, 102 nm. Indices *i* increase from the barbed end to the pointed end. We chose this form for the capture force as while a captured actin filament would have a tendency to move towards the center of the bound Myo2 cluster, it is presumably not realistic for all the filaments bound to a cluster to be forced to occupy an infinitesimally small region at center of the cluster. Thus, we set the capture force to zero if the filament is located sufficiently close to the center of the cluster, and increase the force according to the equation above depending on its distance from the center. See the subsubsection *Myosin unbinding thresholds* for details on how the force constant $k_{\mathrm{Myo2}}$ is set. The formula for Myp2-actin binding force can be obtained from the equation above by replacing $k_{\mathrm{Myo2}}$ with $k_{\mathrm{Myp2}}$ and $d_{Myo2}$ with $d_{Myp2}$ that equals 50 nm.  According to Newton's Third Law, actin subunit *i* exerts an equal and opposite force of $-\vec{f}_{i,\alpha}^{\mathrm{cap}}$ on Myo2 or Myp2 labelled by α.

Myosin unbinding thresholds. Here, we explain the procedure to obtain the values of $k_{\mathrm{Myo2}}$ and $k_{\mathrm{Myp2}}$, the spring constants of the capture forces of the respective myosins. First, we define the unbinding threshold forces of Myo2 and Myp2, $f_{\mathrm{unbind}}^{\mathrm{Myo2}}$ and $f_{\mathrm{unbind}}^{\mathrm{Myp2}}$ as the force attained at the edge of the capture zone i.e. 51 nm from Myo2 capture zone center or 100 nm from Myp2 capture zone center. We obtained these thresholds as best-fit parameters by comparing ring cross sections between simulation and experiment. We measured ring thickness, defined as the spread of 98% of actin segments along the direction perpendicular to the membrane (Figure 4G & 6A), and Myo2-Myp2 separation, defined as the difference of the medians of the distances between Myo2/Myp2 and the plasma membrane, as a function of these two parameters, and saw that when both unbinding thresholds equaled the best-fit values, the ring thickness and Myo2-Myp2 separation matched previous experimental measurements, Figures 4G, 6A, and S4 (Laplante et al., 2016; McDonald et al., 2017).

For simulations of rings in *myo2-E1* cells, the Myo2 unbinding threshold force was lowered to 12 pN so that 1-2 straight bridges of actin filaments and Myp2 detached from the membrane, and that the lengths of these bridges did not exceed roughly a third of the length of the ring. This reproduces experimental observations in *myo2-E1* cells, where ~2 bridges spanning no larger than a third of the ring perimeter were seen in a constricting ring (Laplante et al., 2015).

Number of Myp2 heads per cluster. To determine the number of heads in each Myp2 cluster, we simulated rings varying this parameter while maintaining the total number of Myp2 heads.

We found that with too few heads (fewer than 16), the ring tension is significantly lowered, Figure S3A. We attributed the drop in tension to the fact that, with fewer heads, the net polarity of filaments bound by a Myp2 cluster fluctuates more widely as the number of bound filaments is lower. When a Myp2 clusters is faced with a net polarity of zero, it does not move and is able to produce maximal tension in the ring. When the net polarity is non-zero, Myp2 clusters move relative to the bound filaments, and the forces exerted are lower due to the force-velocity relation. In simulations with Myp2 clusters with too many heads (greater than 16), ring tension was lowered, and the fraction of actin in whiskers was increased substantially, Figure S3B. Only simulations with 16 Myp2 heads per cluster produced ring tensions consistent with our experimental measurements, Figure S3A.

Myosin pulling forces and the force-velocity relation. Each Myo2 or Myp2 cluster applies a pulling force to captured actin segments, with the force acting along the filament, towards the pointed end (see the subsection *Binding of myosin to actin* for details of myosin capture). The magnitude of this force depends on the number of filaments interacting with the cluster, and the relative velocity between each actin filament and myosin, given by the force-velocity relation. We implemented a linear force-velocity relation for simplicity. Below a critical number of captured actin filaments $n^*$, each filament experiences a maximum force of $f_{\text{filament}}$ from a cluster. At the critical number $n^*$, the maximum force exerted on the filaments $n^* f_{\text{filament}}$ equals the maximum force that can be developed by a cluster, which is $16 f_{\text{Myo2/Myp2}}^{\text{stall}}$ as there are only 16 heads per cluster. Above the critical number $n^*$, the maximal myosin force $16 f_{\text{Myo2/Myp2}}^{\text{stall}}$ is distributed equally amongst the filaments. Thus, the myosin force that Myo2 α exerts on actin segment *i* is

$$\vec{f}_{\text{Myo2}}^{i,\alpha} = \begin{cases} f_{\text{filament}} \left[ 1 - \dfrac{(\vec{v}_i - \vec{v}_\alpha) \cdot \hat{\mathrm{t}}_i}{v_{\text{myo}}^0} \right] \hat{\mathrm{t}}_i & n_\alpha < n^* \\ f_{\text{Myo2}}^{\text{stall}} \dfrac{16}{n_\alpha} \left[ 1 - \dfrac{(\vec{v}_i - \vec{v}_\alpha) \cdot \hat{\mathrm{t}}_i}{v_{\text{myo}}^0} \right] \hat{\mathrm{t}}_i & n_\alpha \geq n^* \end{cases}$$

where $\vec{v}_i$ and $\vec{v}_\alpha$ represent the velocities of the actin subunit and myosin II cluster, respectively; $v_{\text{myo}}^0$ is the load-free velocity of myosin II; $n_\alpha$ is the total number of actin filaments captured by Myo2 α; $\hat{t}_i = (\vec{r}_{i+1} - \vec{r}_i)/|\vec{r}_{i+1} - \vec{r}_i|$ is the unit tangent vector pointing towards the pointed end, i.e. pointing from subunit *i* towards the next subunit *i+1*. The formula for Myp2 force $\vec{f}_{\text{Myp2}}^{i,\alpha}$ can be obtained from the equation above by replacing the Myo2 parameters with the corresponding Myp2 parameters. According to Newton's Third Law, actin segment *i* exerts $-\vec{f}_{\text{Myo2}}^{i,\alpha}$ on Myo2 or $-\vec{f}_{\text{Myp2}}^{i,\alpha}$ Myp2 α.

We explain here how we obtained the maximum force exerted on a filament by a node $f_{\text{filament}}$ by comparison with experiment. In a previous experimental study, diffusivity of newly formed nodes $D$ were measured prior to ring assembly and a membrane anchor drag coefficient γ was inferred using the Einstein relation $D = k_B T/\gamma$ (Vavylonis et al., 2008). During ring assembly, the nodes were seen to move ballistically with mean velocity $v$. Thus, the pulling force acting on each node was $\gamma v = 4$ pN. It was later shown using FPALM super-resolution methods that each spot visualized using confocal microscopy during ring assembly was actually two nodes in close proximity (Laplante et al., 2016). Thus, the drag coefficient γ measured was for one node, whereas the velocity measurement $v$ was for 2 nodes in close proximity. Thus, the force on each node was earlier underestimated by a factor of 2 as the drag coefficient was underestimated by a factor of 2. Thus, the pulling force acting on each node is 8 pN. As during assembly node density is very low, on average one node-attached filament is pulled by only one other node. Hence, the 8 pN is a measurement of the force per filament exerted by a cluster $f_{\text{filament}}$ when very few actin filaments are interacting with it (i.e. when $n_\alpha < n^*$).

Anchor drag forces on nodes. Drag forces from the membrane resist the motion of membrane anchored nodes. On the $i^{\text{th}}$ node, the force is

$$\vec{f}^{\,i}_{\text{anch}} = \gamma_{\text{anch}} \vec{v}_i$$

where $\gamma_{\text{anch}}$ is the viscous drag coefficient of an anchor in the membrane and $\vec{v}_i$ is the velocity of the node.

Turnover of components. (i) Nodes bind the membrane in a zone of fixed width with equal probability per unit area, and unbind (are removed from the simulation) stochastically, characterized by a mean lifetime. Nodes entering the ring were only introduced at positions where they would not overlap with existing nodes. (ii) Formins stochastically bind nodes. These rules are described in the subsection *Formin binding and unbinding rates*. For simplicity, we assume that formins do not unbind from nodes; instead, they are removed from the simulation when nodes that they bind to disassociate from the ring. (iii) Formins nucleate an actin filament immediately upon binding a node, i.e. each formin dimer is immediately associated with a 10 nm-long actin "rod". The formin dimer occupies one end of this "rod". The initial orientation of the actin rod is randomly chosen with two constraints. Firstly, the radial component of the tangent vector to the rod must be inward. Secondly, the angle between ring and the tangent vector to the rod must be less than a fixed value, $\theta_{\text{max}}$ (Table S1). All orientations satisfying these constraints are chosen with equal probability. The overall ring structure was not affected by increasing the maximum angle $\theta_{\text{max}}$ or by removing this constraint entirely, though the whisker fraction did increase with larger $\theta_{\text{max}}$ values. (iv) Formins continuously grow actin filaments at rate $v_{\text{pol}}$, i.e. the first actin segment is elongated at this rate. When the rod exceeds 110 nm, it is broken into two by a hinge at a location 10 nm from the formin, i.e. a new rod is added. (v) Actin filaments are stochastically severed, mediated by cofilin. We assume homogeneous severing rate (per unit length). When severed at a location, all actin subunits from that location to the pointed end are removed from the simulation. (vi) Actin filaments are removed from the simulation when the formin dimers they are associated with are removed. (vii) Myp2 clusters stochastically bind to existing actin filaments, i.e. new Myp2 clusters appear in the simulation with equal probability per unit volume as long as an actin filament is within reach. Myp2 clusters unbind stochastically after a mean time . (viii) α-actinin crosslinks stochastically bind pairs of actin subunits within reach (separated by less than the α-actinin length 30 nm), and unbind stochastically after a mean time or when overstretched (length > 50 nm).

Determination of turnover parameters. (i) Unbinding rates of cytokinesis nodes and Myp2 clusters $k^{\mathrm{off}}_{\mathrm{Myo2/Myp2}}$ are directly taken from experiment (Table S1). As no time-course of $k^{\mathrm{off}}_{\mathrm{Myo2/Myp2}}$ is available experimentally, we assume constant $k^{\mathrm{off}}_{\mathrm{Myo2/Myp2}}$ throughout constriction. We averaged off-rates of myosin light chain Cdc4 and formin Cdc12 that were previously measured to obtain the off-rate of Myo2 $k^{\mathrm{off}}_{\mathrm{Myo2}}$ (Table S1). (ii) Binding rates of cytokinesis nodes and Myp2. Densities of Myo2 and Myp2, $\rho_{\mathrm{Myo2/Myp2}}$, were measured by experiment (Figure 3 in (Wu and Pollard, 2005)). We use these and the unbinding rates to set the binding rates $r_{\mathrm{Myo2/Myp2}}$ of cytokinesis nodes and Myp2: $r_{\mathrm{Myo2/Myp2}} = k^{\mathrm{off}}_{\mathrm{Myo2/Myp2}}\rho_{\mathrm{Myo2/Myp2}}$. At four different time points, $\rho_{\mathrm{Myo2/Myp2}}$ has been measured, and we use an interpolating spline fit to these time points to estimate $\rho_{\mathrm{Myo2/Myp2}}$ at any given time during constriction. Using time-dependent $\rho_{\mathrm{Myo2/Myp2}}$ and constant $k^{\mathrm{off}}_{\mathrm{Myo2/Myp2}}$ we obtain the time-course of $r_{\mathrm{Myo2/Myp2}}$. (iii) For binding rates of formin Cdc12p, see the subsection *Formin binding and unbinding rates* below. (iv) Actin polymerization rate $v_{\mathrm{pol}}$ and actin severing rate $r_{\mathrm{sev}}$. These two are set as best-fit parameters by fitting model-predicted mean actin filament length $\bar{l}$ and actin turnover time $\tau_{\mathrm{act}}$ to experiment, where $\tau_{\mathrm{act}}$ is defined as the time during which 90% of actin rods turn over. The experimentally measured time-course of $\bar{l}_{\mathrm{exp}}$ is obtained from (Courtemanche et al., 2016) as the ratio of the total actin filament length and number of formin Cdc12 dimers. The experimental value of $\tau^{\mathrm{exp}}_{\mathrm{act}}$ is taken to be 55 s because treatment of a high dose of Latrunculin A resulted in ring disintegration in 55 s in fission yeast (Yonetani et al., 2008). The model-predicted actin filament length $\bar{l}$ and actin turnover time $\tau_{\mathrm{act}}$ were calculated using the partial differential equation in the subsection *Derivation of the steady-state actin length probability distribution* below (please see that subsection as well).

Formin binding and unbinding rates. Here, we describe the kinetics of formin binding and unbinding from the ring. All formin dimers are bound to membrane-anchored nodes in the simulation. When a node is introduced into the ring, it has no formins. It can bind up to two formin dimers during its lifetime. Formins never unbind from their host nodes; these formins are removed from the simulation only when the host node is removed.

At each timestep, there exist a certain number of “zero”, “one”, and “two” nodes in the simulation, labelled $n_0, n_1,$ & $n_2$ respectively; the labels refer to the number of formins bound to the nodes. The following system of equations are evolved in time for an interval equal to one timestep of the simulation, to determine these numbers at the next timestep:

$$\dot{n}_0 = -k_{01} n_0$$
$$\dot{n}_1 = k_{01} n_0 - k_{12} n_1$$
$$\dot{n}_2 = k_{12} n_1$$

Here $k_{01}$ and $k_{12}$ are the rates of conversion of “zero” into “one”, and “one” into “two” nodes respectively. From these numbers, we obtain the probabilities of a “zero” node converting into a “one” node, and a “one” node converting into a “two” node. Then, we chose nodes at random and increased the number of bound formins on them suitably according to these probabilities.

We chose $k_{01} = 100\, k_{\mathrm{Myo2}}^{\mathrm{off}}$ and $k_{12} = 0.0012\, k_{01}$. These were chosen to set a mean of 1.1 formin dimers per node, which is the value at constriction onset (Table S1), and to set the ratio of numbers of “zero” to “one” nodes at 1%. We chose this ratio as we reasoned that formins would bind very rapidly to a node without any formins. The mean number of formin dimers per node remains close to ~1.1 throughout constriction, so we did not vary these rates throughout constriction.

Initial condition. We describe here the initial condition used in all simulations presented here except those shown in Figure 4D and S6 (in which the ring self-assembles from an initially bare membrane). The ring is equilibrated for 90 s before constriction onset. The values of the parameters in the simulation were either immediately set to their respective values at the onset of constriction, or changed to these values during equilibration (Table S1).

The initial diameter of the ring is 3.7 μm (Laplante et al., 2015). Myo2 nodes are randomly placed along the ring, laterally dispersed in a band of width 200 nm. Initially, there are an average of 0.9 formin Cdc12 dimers per Myo2 node. These are distributed randomly among these nodes, with the rule that each node can contain at most 1 Cdc12 dimer. Each formin dimer caps an actin filament whose length is chosen from the actin length probability distribution given below. This is the steady-state filament length distribution that would have resulted from the processes of filament growth, severing, and turnover of nodes,

$$f_{\mathrm{ss}}(l) = \left[\frac{k_{\mathrm{off}}^{\mathrm{for}} + l\, r_{\mathrm{sev}}}{v_{\mathrm{pol}}}\right] \exp\left[-\frac{l\left(2k_{\mathrm{off}}^{\mathrm{for}} + l\, r_{\mathrm{sev}}\right)}{2v_{\mathrm{pol}}}\right]$$

Here, $f_{\mathrm{ss}}(l)$ is the probability density function of actin filament length $l$, $v_{\mathrm{pol}}$ the polymerization rate, $r_{\mathrm{sev}}$ the cofilin-mediated severing rate and $k_{\mathrm{off}}^{\mathrm{for}} = k_{\mathrm{Myo2}}^{\mathrm{off}}$ the formin off rate because formins unbind the ring with the Myo2 nodes that they are associated with, and do not unbind the ring in any other way. See the subsection *Derivation of the steady-state actin length probability distribution* below for the derivation. Actin filaments lie parallel to the ring (in circular arcs following the curved contour) and have clockwise or counter-clockwise polarity with equal probability. A total of 145 Myp2 clusters are randomly placed along the ring, in a cylindrical band 6 nm further away from the membrane than Myo2.In order to demonstrate the capacity of the ring to self-assemble, we also performed simulations in which the ring built itself up from almost nothing, Figure 4G. In these simulations, 1 node containing no formin is introduced into the ring. Additional nodes, formin Cdc12, actin filaments, and Myp2 are then introduced into the ring according to the turnover rules described above, with all parameters set to their respective values at constriction onset.

Derivation of the steady-state actin length probability distribution. To compute $f_{\mathrm{ss}}(l)$, we denote the number of filaments in the ring with length between $l$ and $l + \Delta l$ as $F(l,t)\Delta l$. The dynamics of the length distribution are

$$\frac{\partial F(l,t)}{\partial t} = r_{\mathrm{nucl}}\delta(l) - v_{\mathrm{pol}}\frac{\partial F(l,t)}{\partial l} - r_{\mathrm{sev}} l F(l,t) + r_{\mathrm{sev}}\int_{l}^{\infty} F(l',t)\mathrm{d}l' - k_{\mathrm{off}}^{\mathrm{for}} F(l,t)$$

where $k_{\mathrm{off}}^{\mathrm{for}}$, $r_{\mathrm{sev}}$ and $v_{\mathrm{pol}}$ are the formin off rate, cofilin-mediated actin severing rate (per filament length) and formin-mediated actin polymerization rate, respectively (Stachowiak et al., 2014). The first term on the right-hand side represents nucleation (with nucleation rate $r_{\mathrm{nucl}}$), the second term polymerization of actin subunits, the third and fourth terms cofilin severing, and the fifth term unbinding of formin from the ring.

Setting $\partial F(l,t)/\partial t = 0$ at steady state, and taking $\partial/\partial l$ of both sides, we have, for $l > 0$:

$$0 = -v_{\mathrm{pol}}\frac{\partial^2 F(l,t)}{\partial l^2} - \left(r_{\mathrm{sev}} l + k_{\mathrm{off}}^{\mathrm{for}}\right)\frac{\partial F(l,t)}{\partial l} - 2 r_{\mathrm{sev}} F(l,t)$$

Solving this equation and normalizing the total probability to unity yields the steady state actin filament length distribution $f_{\mathrm{ss}}(l)$ as shown in the previous subsection, with $\int_0^\infty f_{\mathrm{ss}}(l)\,\mathrm{d}l = 1$, and the mean actin filament length $\langle l \rangle_{f_{\mathrm{ss}}} = \int_0^\infty l f_{\mathrm{ss}}(l)\,\mathrm{d}l = 2.5\ \mu$m.

Running the simulation. We adapted the computation scheme developed in (Platt, 1989). At every time step, the system is described by component positions $\{q_i\}$, i.e. the x, y and z coordinates of all N particles in the system, where i = 1, 2, 3, …, 3N. We use first-order dynamics i.e. forces due to interactions between particles, and constraint forces are resisted by drag forces. Non-constraint forces $\{F_i\}$ are calculated from the force laws as described in previous subsections. Here $\{F_i\}$ do not depend on velocities. Velocity-dependent forces, i.e. myosin forces that obey a linear force-velocity relation, are described as the sum of a velocity-independent part (absorbed into $\{F_i\}$) and the product of velocity with an anisotropic drag coefficient matrix (absorbed into the drag coefficient matrix $\gamma_{ij}$, see below). At each timestep, a system of equations is solved to obtain constraint forces and component velocities simultaneously.

The system is subject to constrains $\{g_i(\{q_j\}) = 0\}$, including (i) Myo2 clusters are constrained at a fixed distance from plasma membrane, i.e. the center of each Myo2 cluster is 94 nm away from the membrane; (ii) formin Cdc12p dimers are bound to nodes at a location between the Myo2 and the membrane, 44 nm from the membrane; (iii) the formin Cdc12p and the Myo2 of a node are anchored to the same point on the membrane; (iv) each actin filament is represented as a chain of 100 nm rods, with the exception that the length of the barbed-end rod is variable (polymerization), i.e. the rod lengths are constrained. These constrains are enforced in a rate-controlled way (with Einstein summation convention):

$$\frac{dg_i}{dt} = \frac{\partial g_i}{\partial t} + \frac{\partial g_i}{\partial x_j}\frac{dx_j}{dt} = -\frac{1}{\tau}g_i$$

Constraint forces, denoted $\{\lambda_i\}$, are calculated together with the velocities of the particles $\{dx_i/dt\}$ by solving the following system of equations.

$$\gamma_{ij}\frac{dx_j}{dt} + \lambda_j\frac{\partial g_j}{\partial x_i} = F_i$$

Recall that the myosin pulling force applied to actin subunits contains a term proportional to the relative velocity of the myosin and actin beads. This velocity dependence is absorbed into the drag matrix $\gamma_{ij}$, while the other, velocity-independent terms are included as contributions to $F_i$. For each actin subunit $\alpha$ subject to pulling force by a Myo2 cluster, the drag matrix is given by

$$\gamma_{ij}^{\alpha} = \gamma_0\delta_{ij} + \sum_m \frac{f_{\mathrm{Myo2}}}{v_{\mathrm{myo}}^0} t_i^{\alpha} t_j^{\alpha},$$

where $\gamma_0$ is the drag of an actin subunit, $\delta_{ij}$ is the Kronecker delta, and $t_i^{\alpha}$ is the $i$ component of the filament's unit tangent vector, and the sum is taken over all Myo2 clusters $m$ pulling the actin subunit. Similarly, for a Myo2 cluster $m$, the drag matrix is given by

$$\gamma_{ij}^{m} = \gamma_{\mathrm{Myo2}}\delta_{ij} - \sum_{\alpha} \frac{f_{\mathrm{Myo2}}}{v_{\mathrm{myo}}^0} t_i^{\alpha} t_j^{\alpha},$$

where the summation is taken over all actin subunits $\alpha$ pulled on by this Myo2 cluster. To obtain the corresponding relation for filaments pulled by Myp2, replace the Myo2 stall force and drag in the two equations above with those of Myp2.

After the velocities are calculated, the ring is evolved using the forward Euler method: the net force on each component is calculated from the positions at some instant, from which the velocities are calculated, and the positions updated by adding the product of the velocities and the timestep 1/30 s. At every time step, components are added to and removed from the simulation according to the turnover rules discussed in the preceding subsections.

**Analytical model of the contractile ring**

Here, we present an analytical calculation of how node velocities and ring tension vary with actin filament length, myosin II density, and the membrane anchor drag coefficient of the nodes in our contractile ring model. We want to illustrate the role played by lateral anchoring of the nodes in tension generation. Our calculation here is a simplified version of the one from our earlier study with a continuum, coarse-grained model of the ring (Thiyagarajan et al., 2017). We have also included the effect of Myp2 here—it was not a part of that study.

Let us first examine the nature of the nodes in the ring. We assume that the nodes and Myp2 clusters are distributed uniformly throughout the ring, consistent with our simulations that show a largely homogeneous ring (Figure 4E). An important organizational feature in our model is that every actin filament was anchored at its barbed end by a formin dimer to a membrane-anchored node. From table S1, we see that there is a mean of ~1.1 formin dimers per node, and hence a mean of about one filament per node.

As the ring is approximately a cylindrical collar whose thickness and width are much smaller than its length, we assume the node-attached filament is oriented along the ring length and can point clockwise or counterclockwise. Thus, the nodes are delineated into two families according to the polarity of the attached filament; let us call these '+' and '-' nodes respectively. The mean velocity of nodes in each family is $\pm v_{\text{node}}$ as each node moves parallel to the force exerted on its associated filament. There is an equal number of either type of node in the ring as the filament orientation is presumably stochastically determined when a new node binds the ring.

Average node velocity. The total force balance on node $i$ is given by

$$\gamma v_i = T_i + F_i$$

where $v_i$ is the velocity of node $i$, $T_i$ is the tension in the associated filament at the barbed end, and $F_i$ is the total pulling force exerted by node $i$ on the actin bundle.

The tension at the barbed end $T_i$ is given by the total pulling force on the associated filament exerted by myosin II,

$$T_i = \frac{l_i \rho_{\mathrm{myo}}}{2n_x} f_{\mathrm{myo}} \left(1 - \frac{v_i + v_{\mathrm{pol}} - v_{\mathrm{node}}}{v_0}\right) + \frac{l_i \rho_{\mathrm{myo}}}{2n_x} f_{\mathrm{myo}} \left(1 - \frac{v_i + v_{\mathrm{pol}} + v_{\mathrm{node}}}{v_0}\right) + \frac{l_i \rho_{\mathrm{myp}}}{n_x} f_{\mathrm{myp}} \left(1 - \frac{v_i + v_{\mathrm{pol}}}{v_0}\right),$$

where $l_i$ is the length of the associated filament, $\rho_{\mathrm{myo}}$ and $\rho_{\mathrm{myp}}$ are the number of Myo2 and Myp2 heads per unit length around the ring, $n_x$ is the number of filaments in the ring cross section, and $v_{\mathrm{pol}}$ is the polymerization rate of actin. Here, the first term comes from pulling forces exerted by nodes with the same polarity as node $i$, the second term comes from nodes of the opposite polarity, and the last term comes from pulling forces from Myp2 clusters. We have assumed that all nodes pulling the filament associated with node $i$ are moving with velocity $\pm v_{\mathrm{node}}$, and that the Myp2 clusters pulling the associated filament are stationary; this is a reasonable approximation, because fluctuations in the node and Myp2 cluster velocity will be averaged out over the ~15 nodes and ~10 Myp2 clusters pulling a typical filament.

The pulling force exerted by node $i$ on the actin bundle is given by

$$F_i = \frac{n_{\mathrm{head}}}{2} f_{\mathrm{myo}} \left(1 - \frac{v_i + v_{\mathrm{pol}} + v_{\mathrm{node}}}{v_0}\right) - \frac{n_{\mathrm{head}}}{2} f_{\mathrm{myo}} \left(1 + \frac{v_i - v_{\mathrm{pol}} - v_{\mathrm{node}}}{v_0}\right) = -\frac{n_{\mathrm{head}} f_{\mathrm{myo}} v_i}{2v_0},$$

where $n_{\mathrm{head}} = 16$ is the number of Myo2 heads per node and the number of Myp2 heads per cluster. Here, the first term arises from forces on nodes of opposite polarity, while the second term arises from nodes of the same polarity.

Inserting the relations derived above for $T_i$ and $F_i$ and solving for the node velocity $v_i$, we find

$$v_i = \frac{l_i}{n_x} \frac{\rho_{\text{myo}} f_{\text{myo}} + \rho_{\text{myp}} f_{\text{myp}}}{\gamma + \frac{n_{\text{head}} f_{\text{myo}}}{v_0} + \frac{l_i \rho_{\text{myo}} f_{\text{myo}}}{n_x v_0} + \frac{l_i \rho_{\text{myp}} f_{\text{myp}}}{n_x v_0}} \left(1 - \frac{v_{\text{pol}}}{v_0}\right).$$

We obtain an approximate formula for the mean speed $v_{\text{node}}$ by replacing $l_i$ in the equation above with the average filament length $\langle l_i \rangle = l_{\text{fil}}$, effectively assuming a monodisperse filament length distribution. This assumption introduces inaccuracies into the analytical model, because the filament length distribution is broad, with a standard deviation similar in magnitude to the mean; however, we still obtain a useful functional form for the average node speed from this procedure,

$$v_{\text{node}} = \frac{l_{\text{fil}}}{n_x} \frac{\rho_{\text{myo}} f_{\text{myo}} + \rho_{\text{myp}} f_{\text{myp}}}{\gamma + \frac{n_{\text{head}} f_{\text{myo}}}{v_0} + \frac{l_{\text{fil}}}{n_x v_0} \left(\rho_{\text{myo}} f_{\text{myo}} + \rho_{\text{myp}} f_{\text{myp}}\right)} \left(1 - \frac{v_{\text{pol}}}{v_0}\right).$$

Thus, if the node anchor drag is very weak, the node velocity is increased. We have used this functional form to fit our simulated node speed vs. node anchor drag coefficient data in Fig. 5I, $v_{\text{node}} = F^*/(\gamma + \gamma^*)$, treating $F^*$ and $\gamma^*$ as fitting parameters.

Total ring tension. Given the above analytical formula for the node velocity, we can estimate the total ring tension starting with the average tension $T_i$ at the barbed end of a filament calculated above. First, we assume that all nodes move at speed $v_{\text{node}}$; in that case, the expression for $T_i$ reduces to

$$T_i = \frac{l_i}{n_x} \left[\rho_{\text{myo}} f_{\text{myo}} + \rho_{\text{myp}} f_{\text{myp}}\right] \left(1 - \frac{v_{\text{node}} + v_{\text{pol}}}{v_0}\right).$$

The ring tension is given by the total tension of the filaments in the cross section. Because the local filament tension increases linearly as a function of distance from the filament's pointed end (see subsection *Analytical prediction of filament and ring tension* below), each filament contributes on average half of its tension at the barbed tend $T_i$ to the ring tension. The probability that a filament with length $l_i$ passes through a randomly chosen cross section of the ring is given by $l_i/L_{\text{ring}}$, where $L_{\text{ring}}$ is the circumference of the ring. The ring tension is therefore given by

$$T = \sum_i \frac{l_i T_i}{2\, L_{\text{ring}}},$$

where the summation is taken over all filaments. Note that the number of filaments in the ring's cross section is given by the ratio of the total length of actin in the ring to the ring circumference, $n_x = \sum_i l_i / L_{\text{ring}}$. Using this to replace the ring circumference $L_{\text{ring}}$, we can therefore write the ring tension as

$$T = \frac{n_x \sum_i l_i T_i}{2 \sum_i l_i} = \frac{n_x \langle l_i T_i \rangle}{2 \langle l_i \rangle},$$

where $\langle \cdot \rangle$ indicates the mean over all filaments in the ring. Inserting our equation for $T_i$, we find

$$T = \frac{\langle l_i^2 \rangle}{2 \langle l_i \rangle} \left[ \rho_{\text{myo}} f_{\text{myo}} + \rho_{\text{myp}} f_{\text{myp}} \right] \left( 1 - \frac{v_{\text{node}} + v_{\text{pol}}}{v_0} \right).$$

Interestingly, the total ring tension is proportional to the second moment of the filament length distribution, $\langle l_i^2 \rangle$. Taking the approximation that all actin filaments are the same length for simplicity, we see that the ring tension is proportional to the filament length.

Lastly, we note that our simulations showed a small amount of actin (~6%) in the cytokinetic ring existed as disordered whiskers, not assembled into the bundle. Typically, these whiskers formed when the pointed end of a filament protruded out from the bundle. These segments of actin are not pulled by myosin II and hence do not contribute to ring tension. To account for this, we multiple the length of all filaments by a factor $1 - w$, where $w$ is the whisker fraction, in the equation for the ring tension,

$$T = \frac{l_{\text{fil}}}{2} \left[ \rho_{\text{myo}} f_{\text{myo}} + \rho_{\text{myp}} f_{\text{myp}} \right] \left( 1 - \frac{v_{\text{node}} + v_{\text{pol}}}{v_0} \right) \frac{(1 - w) \langle l_i^2 \rangle}{l_{\text{fil}}^2}.$$

Aside from corrections due to relative motion between actin and myosin, whiskering, and the width of the actin filament length distribution, note that that the ring tension obeys the scaling relation $T \sim l_{\text{fil}} \rho_{\text{myo}} f_{\text{myo}}$, where here we use the subscript "myo" to refer to both myosin II isoforms.

<u>Mechanisms of tension production</u>. Recall that in the above derivation of the average node velocity, we found contributions to the pulling tension in a filament from three sources: (i) nodes of the same polarity, (ii) nodes of opposite polarity, and (iii) Myp2 clusters. Indeed, the total tension at the barbed end of filament $i$ can be written

$$T_i = \frac{l_i \rho_{\text{myo}}}{2n_x} f_{\text{myo}} \left(1 - \frac{v_{\text{pol}}}{v_0}\right) + \frac{l_i \rho_{\text{myo}}}{2n_x} f_{\text{myo}} \left(1 - \frac{v_{\text{pol}} + 2v_{\text{node}}}{v_0}\right) + \frac{l_i \rho_{\text{myp}}}{n_x} f_{\text{myp}} \left(1 - \frac{v_{\text{node}} + v_{\text{pol}}}{v_0}\right).$$

Preserving the separation of these terms, the total ring tension is given by

$$T = \frac{n_x \langle l_i T_i \rangle}{2 \langle l_i \rangle} = \frac{\langle l_i^2 \rangle}{2 \langle l_i \rangle} \left[\frac{\rho_{\text{myo}}}{2} f_{\text{myo}} \left(1 - \frac{v_{\text{pol}}}{v_0}\right) + \frac{\rho_{\text{myo}}}{2} f_{\text{myo}} \left(1 - \frac{v_{\text{pol}} + 2v_{\text{node}}}{v_0}\right) + \rho_{\text{myp}} f_{\text{myp}} \left(1 - \frac{v_{\text{node}} + v_{\text{pol}}}{v_0}\right)\right].$$

Inserting the equation for $v_{\text{node}}$, and noting that $n_x = l_{\text{fil}} \rho_{\text{myo}} / n_{\text{head}}$, we find

$$T = \frac{\langle l_i^2 \rangle}{2 \langle l_i \rangle} \left(1 - \frac{v_{\text{pol}}}{v_0}\right) \left[\frac{\rho_{\text{myo}}}{2} f_{\text{myo}} + \frac{\rho_{\text{myo}}}{2} f_{\text{myo}} \left(\frac{\gamma v_0 - \frac{l_{\text{fil}}}{n_x} \rho_{\text{myp}} f_{\text{myp}}}{\gamma v_0 + \frac{l_{\text{fil}}}{n_x} \left(2\rho_{\text{myo}} f_{\text{myo}} + \rho_{\text{myp}} f_{\text{myp}}\right)}\right) + \rho_{\text{myp}} f_{\text{myp}} \left(\frac{\gamma v_0 + \frac{l_{\text{fil}}}{n_x} \rho_{\text{myo}} f_{\text{myo}}}{\gamma v_0 + \frac{l_{\text{fil}}}{n_x} \left(2\, \rho_{\text{myo}} f_{\text{myo}} + \rho_{\text{myp}} f_{\text{myp}}\right)}\right)\right].$$

We identify the first term in square brackets as the contribution from pulling forces exerted between nodes with the same polarity,

$$T_{\text{fixed}} = \frac{\langle l_i^2 \rangle}{4 \langle l_i \rangle} \left(1 - \frac{v_{\text{pol}}}{v_0}\right) \rho_{\text{myo}} f_{\text{myo}}.$$

Each family of nodes forms a chain around the ring, circling the ring with average velocity $v_{\text{node}}$. Since there is little relative motion between nodes of the same family, this mechanism of tension production is termed *fixed filament*.

The other two terms in square brackets in the ring tension formula above arise from pulling forces between nodes of opposite polarity, and from pulling forces exerted by Myp2 clusters. Nodes of opposite polarity have a large relative speed $2v_{\text{node}}$, while nodes of either polarity move at speed $v_{\text{node}}$ relative to Myp2 clusters, on average. This mechanism of ring tension production is therefore called *sliding filament*; the sliding filament contribution to the ring tension is given by

$$T_{\text{sliding}} = \frac{\langle l_i^2 \rangle}{2\,\langle l_i \rangle}\left(1 - \frac{v_{\text{pol}}}{v_0}\right)\left[\frac{\rho_{\text{myo}}}{2} f_{\text{myo}}\left(\frac{\gamma v_0 - \frac{l_{\text{fil}}}{n_x}\rho_{\text{myp}} f_{\text{myp}}}{\gamma v_0 + \frac{l_{\text{fil}}}{n_x}\left(2\rho_{\text{myo}} f_{\text{myo}} + \rho_{\text{myp}} f_{\text{myp}}\right)}\right)\right.$$

$$\left. + \rho_{\text{myp}} f_{\text{myp}}\left(\frac{\gamma v_0 + \frac{l_{\text{fil}}}{n_x}\rho_{\text{myo}} f_{\text{myo}}}{\gamma v_0 + \frac{l_{\text{fil}}}{n_x}\left(2\,\rho_{\text{myo}} f_{\text{myo}} + \rho_{\text{myp}} f_{\text{myp}}\right)}\right)\right].$$

Note that the sliding filament contribution depends on the node anchor drag coefficient $\gamma$, while the fixed filament contribution does not. When the node anchor drag coefficient is very small, this mechanism contributes a negligible amount to the overall ring tension, as seen in Fig. 5I.

Analytical prediction of filament and ring tension. Here, we calculate an analytical expression for the typical tension profile along the length of one actin filament in our simulated ring, and then use it to calculate the overall ring tension. We assume a homogeneous ring with a uniform density of both anchored nodes and unanchored Myp2 clusters, consistent with our simulations that show a largely homogeneous ring (Figure 4E). Each node in our simulation hosts barbed-end anchored actin filaments and Myo2 molecules. Thus, the distribution of Myo2 and Myp2 heads along an actin filament is uniform. Thus, the force per unit length $f_{\text{fil}}$ felt by an actin filament is given by

$$f_{\text{fil}} = f_{\text{Myo2}}\rho_{\text{Myo2}}^{\text{flt}} + f_{\text{Myp2}}\rho_{\text{Myp2}}^{\text{flt}}.$$

Here, $f_{\text{Myo2}}$ and $\rho_{\text{Myo2}}^{\text{flt}}$ are the force per head of Myo2 on the actin filament, and the density of Myo2 heads along the filament respectively; the subscript Myp2 has a similar meaning. A positive force here means that the force is directed away from the barbed end. Because the filament is elongating at a rate $v_{\text{pol}}$, the force per Myo2 or Myp2 head is given by $f_{\text{Myo2}} = f_{\text{Myo2}}^{\text{stall}}(1 - v_{\text{pol}}/v_{\text{myo}}),\ f_{\text{Myp2}} = f_{\text{Myo2}}^{\text{stall}}(1 - v_{\text{pol}}/v_{\text{myo}})$, where the corresponding stall forces are $f_{\text{Myo2}}^{\text{stall}}$ and $f_{\text{Myp2}}^{\text{stall}}$.

The expression for $f_{\text{fil}}$ above can be written in a more convenient form as

$$f_{\text{fil}} = c_{\text{bound}}\bar{f}_{\text{myo}},$$

where $c_{\text{bound}} = \rho_{\text{Myo2}}^{\text{flt}} + \rho_{\text{Myp2}}^{\text{flt}}$ is the total density of myosin along the filament, and $\bar{f}_{\text{myo}}$ is the weighted average of the two myosin forces, given by

$$\bar{f}_{\text{myo}} = \alpha f_{\text{Myo2}} + (1-\alpha) f_{\text{Myp2}}.$$

Here, $\alpha = \rho_{\text{Myo2}}^{\text{flt}} / c_{\text{bound}}$ is the fraction of myosin II molecules that are Myo2. The filament tension is given by $dT_{\text{fil}}(x)/dx = f_{\text{fil}}$ where $x$ is the coordinate along the filament length and $x = 0$ is the pointed end, where $T_{\text{fil}} = 0$. Thus, filament tension is

$$T_{\text{fil}}(x) = c_{\text{bound}} \bar{f}_{\text{myo}} x.$$

This equation explains the dependence of the ring tension on actin filament length. Since the local filament tension is proportional to distance from the filament's barbed end, the mean filament tension scales as the total length of the filament. Thus, longer filaments produce proportionally increased ring tension.

We saw a good agreement between this analytical prediction and the tension along a filament measured in our simulation (Figure 5B). The values of the parameters $c_{\text{bound}}, v_{\text{pol}}, \rho_{\text{Myo2}}^{\text{flt}}$, and $\rho_{\text{Myp2}}^{\text{flt}}$ used there were measured from the simulation at 0 min after constriction onset.

To calculate the ring tension, we first observe that as the filament tension profile is linear, the mean tension per filament is $c_{\text{bound}} \bar{f}_{\text{myo}} l_{\text{fil}}/2$ where $l_{\text{fil}}$ is the mean actin filament length. To get the total ring tension, we just replace $c_{\text{bound}}$ which is the total density of myosin heads along the filament with $c_{\text{bound,ring}} = n\rho_{\text{Myo2}} + m\rho_{\text{Myp2}}$ which is the total density of myosin heads along the ring, where $\rho_{\text{Myo2}}$ and $\rho_{\text{Myp2}}$ are the densities of nodes and Myp2 clusters respectively, and $n = m = 16$ is the numbers of heads per Myo2 node or Myp2 cluster. This procedure is equivalent to multiplying the mean tension per filament by the number of filaments in the cross-section $n_{\text{fil}}$, as $c_{\text{bound,ring}}/c_{\text{bound}} = n_{\text{fil}}$. Thus, ring tension is

$$T = c_{\text{bound,ring}} \bar{f}_{\text{myo}} l_{\text{fil}}/2.$$

Dependence of ring tension on component concentrations: In the previous subsubsection, we saw that ring tension is proportional to myosin density and the mean length of filaments i.e.

$T \sim \rho_{\mathrm{myo}} l_{\mathrm{fil}}$, where we now use $\rho_{\mathrm{myo}}$ for the mean density of myosin II in the ring, with each isoform's concentration weighted by its respective stall force. The overall ring tension T is the average of the total tension of all the filaments in the ring's cross section, $T = n_{\mathrm{fil}} T_{\mathrm{fil}}$. Using the fact that the number of filaments in the cross section is given by $n_{\mathrm{fil}} = \mathrm{L_{act}}/\mathrm{L_{ring}}$, where $L_{\mathrm{act}}$ is the total length of F-actin in the ring, we find that $T_{fil} = T/n_{fil} = \rho\, l_{act}\, L_{ring}/L_{act} = L_{ring}\ \rho/N_{fil}$, where $N_{fil}$ is the total number of actin filaments in the ring. Thus the filament tension is proportional to the density of myosin II in the ring at fixed filament number.

Similarly, we see that $T_{fil} = T/n_{fil} = T\, L_{ring}/L_{act} = T\, L_{ring}/(N_{fil}\, l_{act}) = L_{ring}/N_{fil}\, T/l_{act}$. Thus the filament tension scales as $T/l_{act}$ at fixed ring length and number of actin filaments.

During maturation, the number of formin Cdc12 dimers is constant (Courtemanche et al., 2016), thus the number of filaments in the ring is constant. Thus, the number of filaments in the cross-section is proportional to mean filament length $l_{\mathrm{fil}}$. As the mean tension per filament $T_{fil}$ is the total ring tension divided by the number of filaments in the cross-section, we get $T_{\mathrm{fil}} \sim T/l_{\mathrm{fil}} \sim \rho_{\mathrm{myo}}$.

### QUANTIFICATION AND STATISTICAL ANALYSIS

Null-hypothesis testing for statistical significance was performed using GraphPad Prism 6.0f for Mac (La Jolla, CT) or MATLAB (MathWorks). Statistical significance for linear fit for membrane tension, and ring tension was established by an F test with null-hypothesis of zero slope. Statistical differences for ring tension measurements between wild type and myosin mutants were determined by the Student's t-test. Statistical parameters for individual experiments or simulation runs are indicated in the figure legends. Comparison of experimental and simulated ring tensions was performed using a $\chi^2$ difference test between the measured and simulated tension profiles in WT cells and the overall average tension in mutant cells, using the variance in the experimental data as the error.

**Table S1. Key parameter values of the ring simulation. Related to Figure 4.**

| Symbol | Meaning | Value at onset of constriction | Legend |
|---|---|---|---|
| | Ring binding zone width | 0.2 μm | **(A)** |
| | Initial ring length | 11.6 μm | **(B)** |
| | Ring constriction rate | 70 nm $min^{-1}$ | **(B)** |
| $\rho_{\text{Cdc12p}}$ | Density of formin Cdc12p dimers along the ring | 20 $\mu m^{-1}$ | **(C)*** |
| $\rho_{\text{Myo2}}$ | Density of Myo2 nodes along the ring | 18 $\mu m^{-1}$ | **(D)*** |
| $\rho_{\text{Myp2}}$ | Density of Myp2 clusters along the ring | 12.5 $\mu m^{-1}$ | **(E)*** |
| $k_{\text{Myo2}}^{\text{off}}$ | Myo2 off rate | 0.0245 $s^{-1}$ | **(F)** |
| $k_{\text{Myp2}}^{\text{off}}$ | Myp2 off rate | 0.026 $s^{-1}$ | **(G)** |
| $f_{\text{Myo2}}^{\text{stall}}$ | Myo2 stall force per head | 1.75 pN | **(H)** |
| $f_{\text{Myp2}}^{\text{stall}}$ | Myp2 stall force per head | 1.0 pN | **(H)** |
| | Major axis of Myo2 capture zone | 132 nm | **(I)** |
| | Minor axes of Myo2 capture zone | 102 nm (both) | **(I)** |
| | Diameter of Myp2 capture zone | 200 nm | **(J)** |
| $\rho_{\text{actinin}}$ | Density of α-actinin dimers along the ring | 25 $\mu m^{-1}$ | **(K)** |
| $k_{\text{actinin}}^{\text{off}}$ | α-actinin off rate | 3.3 $s^{-1}$ | **(L)** |
| $r_{\text{sev}}$ | Cofilin-mediated severing rate per unit length on actin filament | 0.93 $\mu m^{-1}$ $min^{-1}$ | **(M)*** |

| | | | |
|---|---|---|---|
| $v_{\text{pol}}$ | Formin-mediated actin polymerization rate | 127 nm s$^{-1}$ | **(M)*** |
| $l_{\text{p}}$ | Actin filament persistence length | 10 μm | **(N)** |
| $\gamma_{\text{anch}}$ | Anchor drag coefficient per node | 500 pN·s μm$^{-1}$ | **(O)** |
| $k_{\text{ex}}^{\text{Myp2-Myp2}}$ | Myp2-Myp2 excluded volume spring constant | 0.32 pN nm$^{-1}$ | |
| $k_{\text{ex}}^{\text{Myo2-Myo2}}$ | Myo2-Myo2 excluded volume spring constant | 0.12 pN nm$^{-1}$ | |
| $k_{\text{ex}}^{\text{Anchor-Myp2}}$ | Myo2-Myo2 excluded volume spring constant | 0.095 pN nm$^{-1}$ | |
| $r_0^{\text{Myp2-Myp2}}$ | Myp2-Myp2 excluded volume cutoff distance | 200 nm | |
| $r_0^{\text{Myo2-Myo2}}$ | Myo2-Myo2 excluded volume cutoff distance | 132 nm | |
| $r_0^{\text{Anchor-Myp2}}$ | Anchor-Myp2 excluded volume cutoff distance | 202 nm | |
| $f_{\text{act}}$ | Actin-Actin excluded volume maximum force | 10 pN | |
| $r_{\text{ex}}$ | Actin-Actin excluded volume distance | 15 nm | |
| $f_{\text{filament}}$ | Actin-limited stall force | 8 pN | **(P)** |
| $\text{f}_{\text{unbind}}^{\text{Myo2}}$ | Myo2-actin unbinding threshold force | 40 pN | **(Q)** |
| $\text{f}_{\text{unbind}}^{\text{Myp2}}$ | Myp2-actin unbinding threshold force | 30 pN | **(Q)** |
| $v_{\text{myo}}^0$ | Load-free Myo2/Myp2 velocity | 0.24 μm s$^{-1}$ | **(R)** |
| $k_{\text{x}}$ | Crosslinker spring constant | 25 pN μm$^{-1}$ | **(S)** |
| $r_{\text{x}}$ | Crosslinker rest length | 30 nm | **(S)** |

| | | | |
|---|---|---|---|
| $\theta_{\max}$ | Maximum angle between new actin filament and ring | 30° | **(T)** |

**Legend:**

* indicates that these parameters were tuned throughout constriction to ensure the time course of component numbers matched previous experimental measurements.

**(A)** We assume that new constriction nodes bind the leading edge of the septum (in-growing cell wall). The septum has width ~ 0.2 μm (Cortes et al., 2007).

**(B)** Measured in (Pelham and Chang, 2002).

**(C)**200 dimers of Cdc12 were measured at constriction onset for a ring with circumference ~10 μm. (Courtemanche et al., 2016). For simplicity we neglect formin For3p that is also present, since For3p grows actin cables (Feierbach and Chang, 2001) and *Δfor3* cells have normal actin levels in the ring and lack observable cytokinesis defects.

**(C)** (Yonetani et al., 2008). Together with the value for $\rho_{\mathrm{Cdc12p}}$, this gives a formin binding rate of $r_{\mathrm{Cdc12p}} = 0.46\ \mu\mathrm{m}^{-1}\cdot\mathrm{s}^{-1}$ at the onset of constriction.

**(D)** Calculated from 2900 Myo2p in a 10 μm long ring at the onset of constriction (Wu and Pollard, 2005), and 8 Myo2 dimers per node (Laplante et al., 2016).

**(E)** Calculated from 2000 Myp2p in a 10 μm long ring at the onset of constriction (Wu and Pollard, 2005), and 16 Myp2 heads per cluster, determined in this study (see STAR Methods).

**(F)** Lifetimes of myosin light chains Cdc4p (Pelham and Chang, 2002) and Rlc1 (Clifford et al., 2008) and the formin Cdc12 (Yonetani et al., 2008) were measured previously using FRAP. We calculated the off-rate from the Rlc1 lifetime and used it for the Myo2 off rate as this was in the middle of the range of experimental values.

**(G)** Consistent with off-rates of $0.022\ \mathrm{s}^{-1}$ obtained using previous FRAP measurements of Myp2 (Takaine et al., 2015).

**(H)** Obtained as best-fit parameters by comparing simulated and experimental tension time courses (Figures 5 and S5) in this work.

**(I)** From the distribution of Myo2 heads measured in FPALM (Laplante et al., 2016).

**(J)** From the apparent size of Myp2 clusters measured in deconvolution microscopy (Takaine et al., 2015).

**(K)** (Wu and Pollard, 2005).

**(L)** (Li et al., 2016).

**(M)** Obtained as best-fit parameters in this work, by comparing actin turnover time and mean filament length from previous measurements (see STAR Methods).

**(N)** (Ott et al., 1993; Riveline et al., 1997).

**(O)** Chosen such that model prediction of node velocity (21 ± 10 nm $s^{-1}$, Figure 5H) agrees with Myo2 node velocity (22 ± 10 nm $s^{-1}$) measured by FPALM (Laplante et al., 2016).

**(P)** (Vavylonis et al., 2008)

**(Q)** Obtained as best-fit parameters in this work, by comparing ring thickness and Myo2-Myp2 radial separation between simulation and experiment (Laplante et al., 2016; McDonald et al., 2017) (see Figure S4 and STAR Methods).

**(R)** Calculated from *in vitro* gliding filament assays in (Stark et al., 2010) (see STAR methods).

**(S)** Estimated from *in vitro* measurements on crosslinked actin bundles (Claessens et al., 2006) and from measurements of α-actinin length using electron microscopy (Meyer and Aebi, 1990).

(**T**) Chosen to minimize the fraction of actin in whiskers. The overall structure of the ring was qualitatively insensitive to increases in this parameter, though the whisker fraction increases with larger maximum angles.

**Table S2. List of *S. pombe* strains used in this study. Related to Figure 1.**

| Strain | Genotype | Reference |
|---|---|---|
| AR581 | *h rlc1-tdTomato-natMX6 sad1-GFP-kanMX6 bgs1Δ::ura4*$^{+}$ *P*$_{bgs1}$$^{+}$*::GFP-bgs1*$^{+}$*:leu1*$^{+}$ *leu1-32 ura4-D18 his3-D1 ade6-M21X* | (Arasada and Pollard, 2014) |
| AR619 | *h Δmyp2::his7*$^{+}$ *rlc1-tdTomato-natMX6 sad1-GFP-kanMX6 bgs1Δ::ura4*$^{+}$ *P*$_{bgs1}$$^{+}$*::GFP-bgs1*$^{+}$*:leu1*$^{+}$ *leu1-32 ura4-D18 his3-D1 ade6-M21X* | (Arasada and Pollard, 2014) |
| CL4 | *h- rlc1-3GFP ade6-M216 his3-D1 leu1-32 ura4-D18* | (Vavylonis et al., 2008) |
| CL55 | *h+ myo2-E1 Rlc1-tdTomato-NatMX6 Sad1-mEGFP-KanMX6* | (Balasubramanian et al., 1998) |
| QC240 | *h- rlc1-tdTomato-natMX6 ade6-M210 leu1-32 ura4-D18 41xGFP-CHD-Leu* | (Chen and Pollard, 2011) |

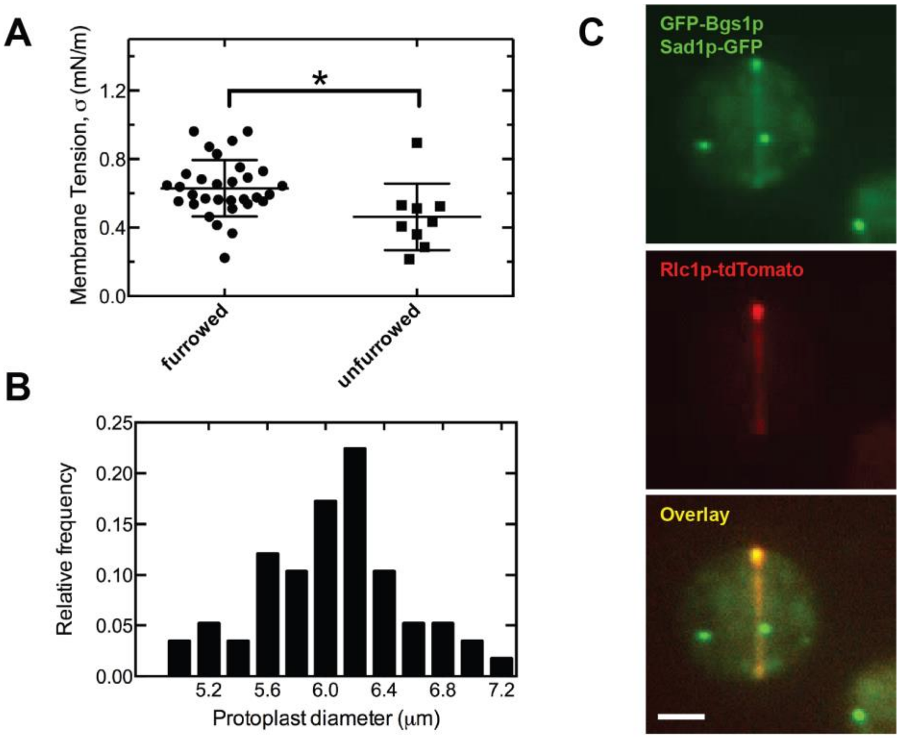

**Figure S1. Furrowed protoplasts are variable in size, have higher membrane tension, and recruit septum synthesis machinery to cytokinetic rings. Related to Figure 2.**

**(A)** Comparison of measured membrane tension in furrowed protoplasts containing rings (same protoplasts as in Figure 2, $n$ = 31) versus nonfurrowed, spherical protoplasts lacking rings ($n$ = 9). Furrowed mitotic protoplasts have membrane tension $\sigma_1 = 0.63 \pm 0.16$ mN m$^{-1}$ (mean ± SD), significantly greater than the unfurrowed interphase protoplast membrane tension $\sigma = 0.46 \pm 0.19$ mN m$^{-1}$ (*$p < 0.05$; two-tailed t-test). Error bars indicate SD. Membrane tensions of unfurrowed protoplasts are not significantly different from those measured previously in (Stachowiak et al., 2014) ($p = 0.18$, two-tailed t-test).

**(B)** Histogram of furrowed protoplast diameters (same protoplasts as in Figure 2, $n$ = 31). Protoplasts vary in size with average diameter $D_{\mathrm{p}} = 5.9 \pm 0.4$ $\mu$m (mean $\pm$ SD).

**(C)** Confocal microscopy images of one protoplast with cytokinetic ring expressing fluorescently-tagged myosin light chain Rlc1p-tdTomato (center), $\beta$-glucan synthase GFP-Bgs1p (top), and spindle pole body marker Sad1p-GFP (top). Rlc1p-tdTomato colocalizes with GFP-Bgs1p (overlay, bottom). Sum intensity projections are displayed. Scale bar = 2 $\mu$m.

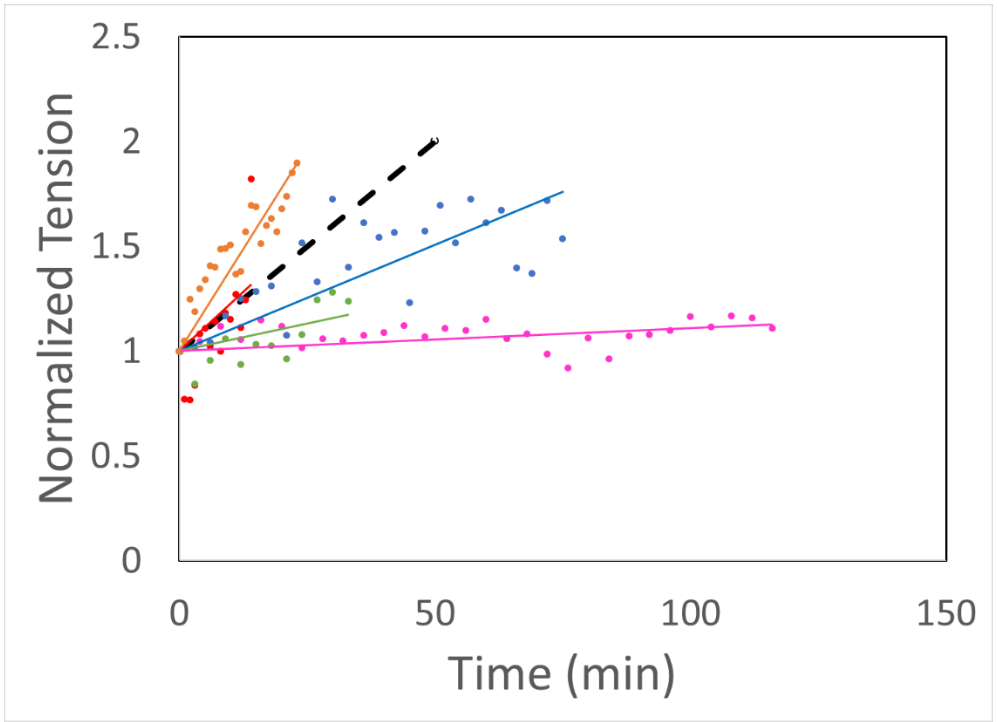

**Figure S2. Cells tensions increased at rates in the range ~0.1 to ~5% per min. Related to Figure 2.**

Contractile ring tension versus time for individual rings as they constrict (solid lines). Each color corresponds to a different cell. Tensions are normalized to the initial value. The dashed line represents the estimated time-dependence of ring tension in cells used for single tension measurements only, based on the tensions measured in Figure 2B and an assumed constriction time in protoplasts of 50 min (Stachowiak et al., 2014).

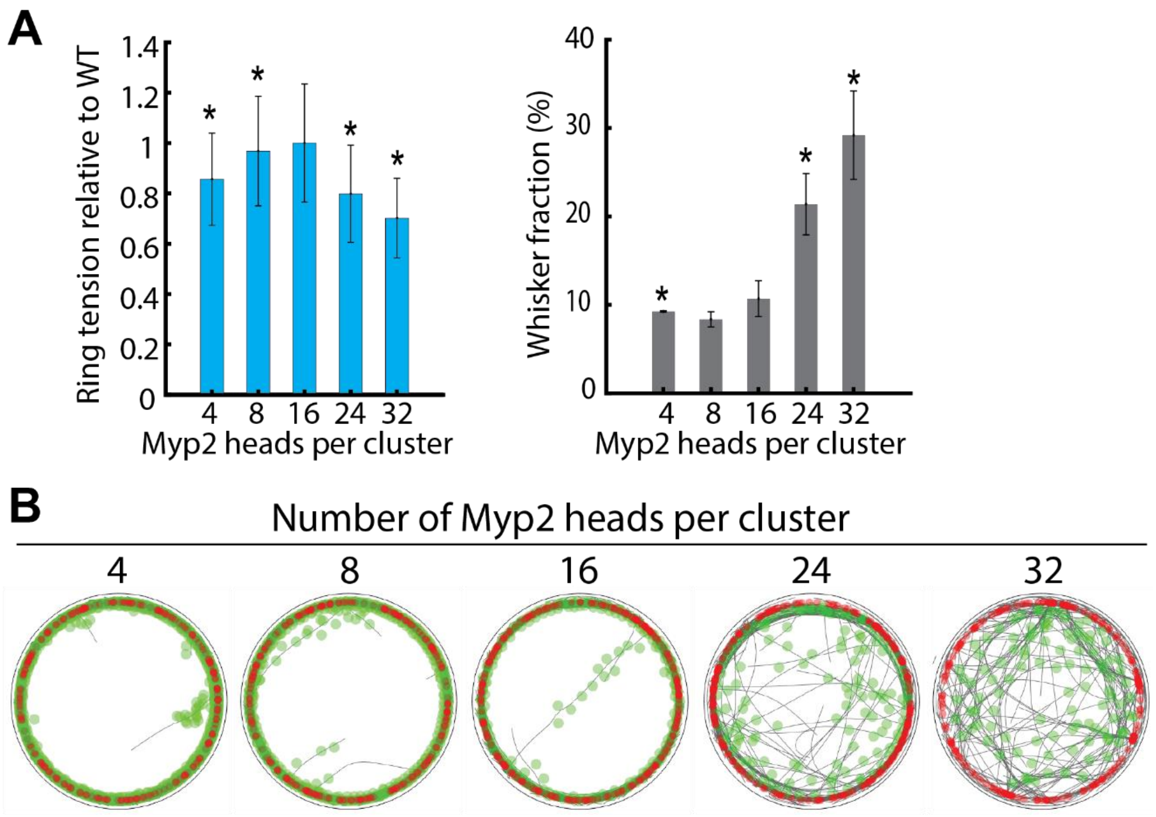

**Figure S3. In simulations the number of heads per Myp2 cluster was chosen to be 16 to best reproduce experimental ring tensions and Myp2 organization. Related to Figure 5.**

**(A)** Simulated ring tensions and whisker fractions with different number of Myp2 heads per cluster, and experimental value (n = 8 for each value of number of Myp2 heads per cluster, and n = 31 for experiment). Simulation parameters as in Table S1 except for the number of Myp2 heads per cluster (as indicated). Error bars represent SD. ns, not significant; ****, $p < 0.05$ by Student's t test. Simulations with greater or fewer than 16 Myp2 heads per cluster produced ring tensions incompatible with experiment.

**(B)** Images of simulated rings 10 min after constriction onset. Simulation parameters as in Table S1 except for the number of Myp2 heads per cluster (as indicated). Myp2 fails to bundle the ring effectively with greater than 16 heads per cluster, causing Myp2 ring fracture, high fractions of actin filaments in whiskers, and Myp2-decorated actin bridges.

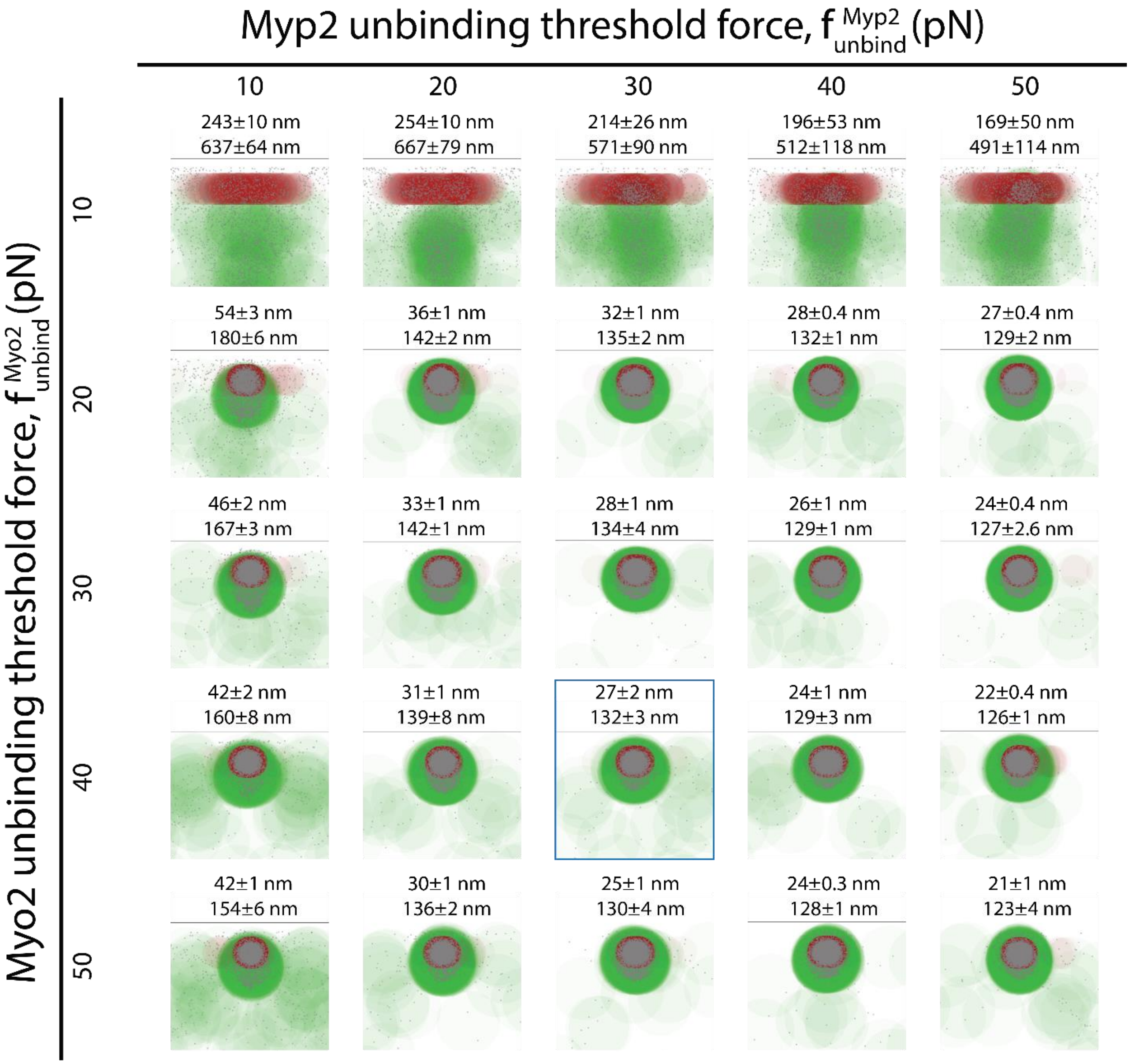

**Figure S4. Integrity of the ring requires sufficiently strong binding of myosin II to actin. Related to Figure 4.**

Each panel shows distribution of Myo2, Myp2 and actin in the cross section of the ring at constriction onset, made using the same procedure used to generate the cross sections in Figure 6Text above each panel has two numbers: the separation of Myo2 and Myp2 (top), calculated as the difference of the medians of the distances between Myo2/Myp2 and the plasma membrane, and the thickness of the ring, defined as the spread of 98% of actin beads in the direction perpendicular to the membrane (bottom). Blue box indicates the parameter values that best reproduce the experimentally measured thickness of the actin bundle, 125 nm (Laplante et al., 2016), and separation between Myo2 and Myp2 heads, 26 nm (McDonald et al., 2017). Values of $f_{\mathrm{unbind}}^{\mathrm{Myo2}}$ less than 30 pN were excluded because these rings formed bridges as the ring radius decreased. Simulation parameters as in Table S1 unless otherwise specified.

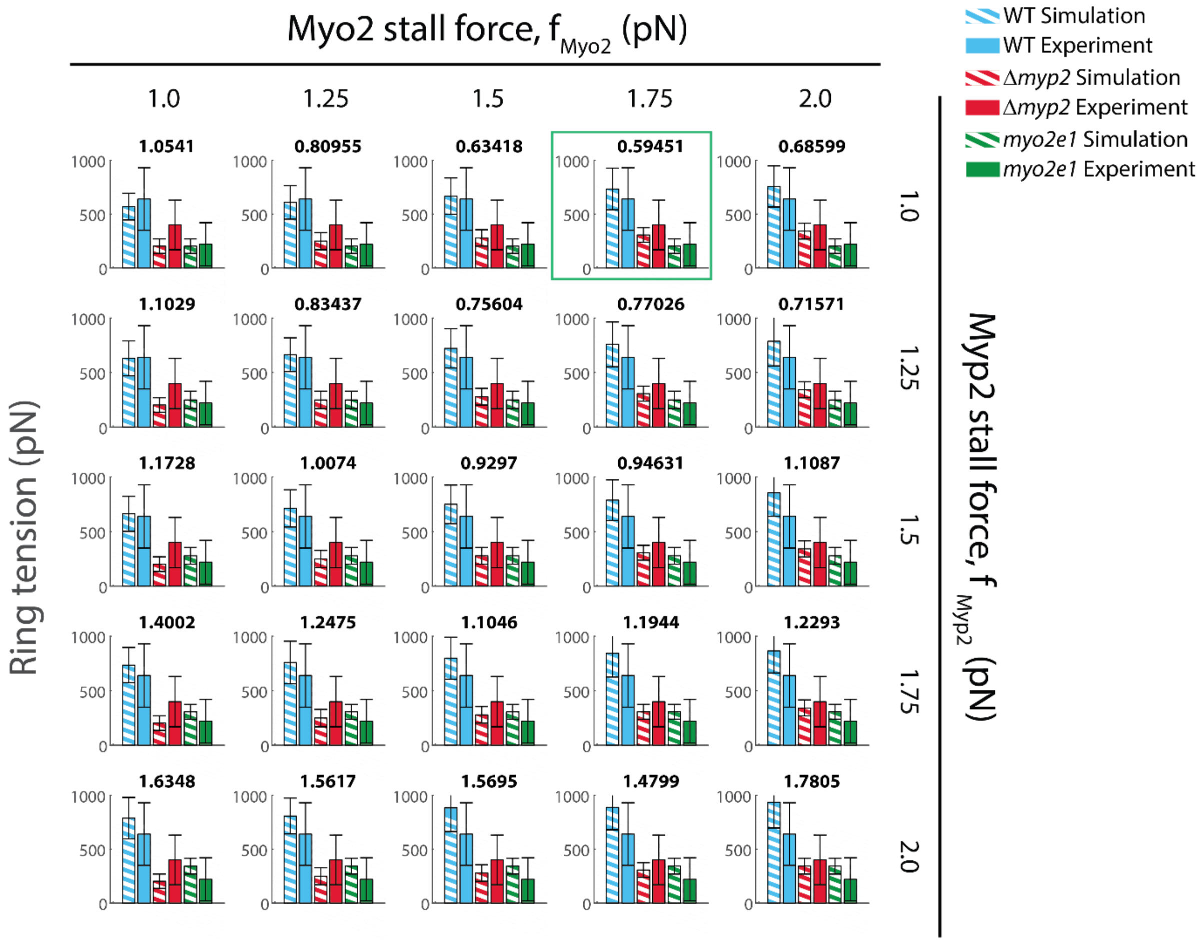

**Figure S5. Results of a scan of Myo2 and Myp2 stall forces per head, $f^{\text{stall}}_{\text{Myo2}}$ and $f^{\text{stall}}_{\text{Myp2}}$. Related to Figure 5.**

Model-predicted mean ring tensions (striped bars, mean ± SD, *n* = 10 simulations for each combination of $f^{\text{stall}}_{\text{Myo2}}$ and $f^{\text{stall}}_{\text{Myp2}}$) were compared to experiment (solid bars, data from Figure 3C). Other simulation parameters as in Table S1. Best-fit parameters $f^{\text{stall}}_{\text{Myo2}} = 1.75$ pN, $f^{\text{stall}}_{\text{Myp2}} = 1.$ pN (green box) minimize the total chi-square statistic (above plots) between the average tension values, using the variance in the experimental data as the error. Lowest chi-square statistic $\chi^2 = 0.59$. See STAR Methods for details.

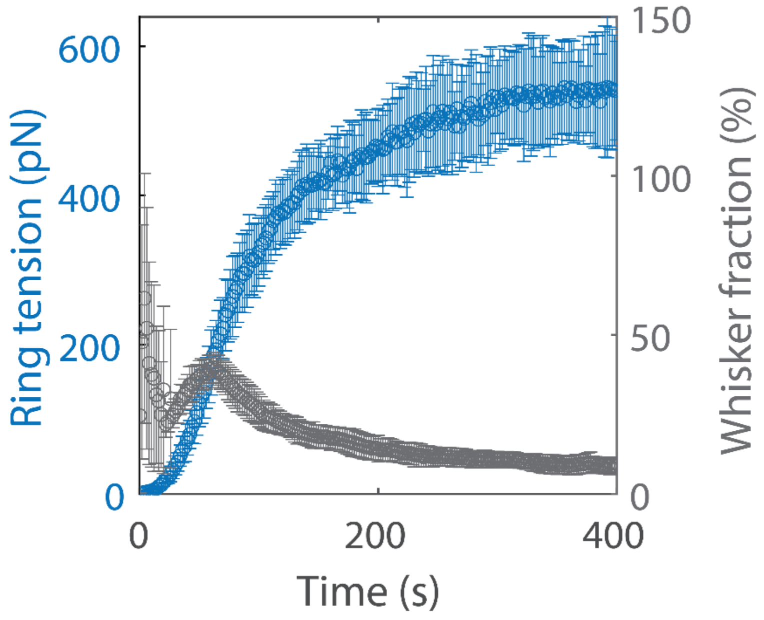

**Figure S6: Rings self-assembled starting from a bare membrane. Related to Figure 4.** Gray: fraction of actin in whiskers in self-assembling rings. Blue: tension in self-assembling rings. After a transient lasting ~3-5 minutes, rings self-assembled into a tense circular actomyosin bundle with few whiskers. Error bars: SD, $n = 30$. The increase in the whisker fraction occurring around 60 seconds is an artifact caused by the small amount of actin that is present in the simulation at this time.

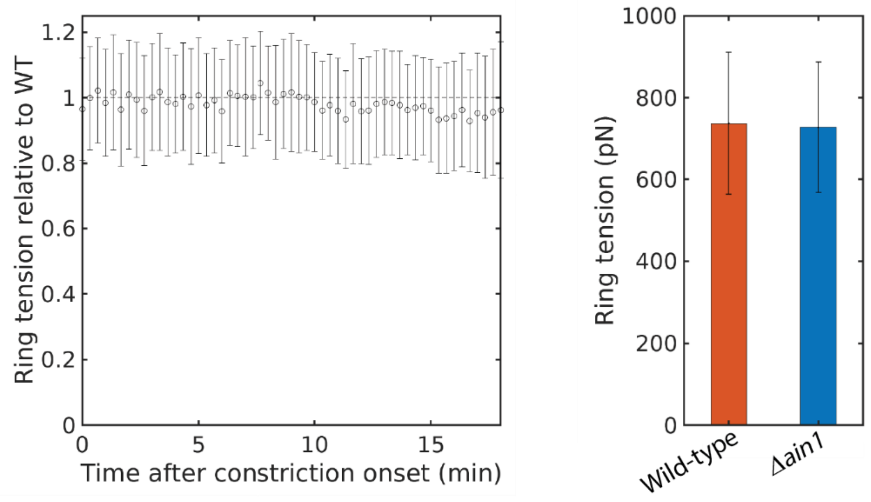

**Figure S7. Removal of α-actinin does not change ring tension significantly.**

Simulated rings with no α-actinin produced tensions that were not significantly different from that of wild-type rings at any point during constriction. Left: ratio of tension in rings without α-actinin to tension in WT rings (n=30). Right: overall average tension in WT rings and those without α-actinin. Error bars: SD.

### STAR METHODS References

Alexander, S.P., and Rieder, C.L. (1991). Chromosome Motion during Attachment to the Vertebrate Spindle - Initial Saltatory-Like Behavior of Chromosomes and Quantitative-Analysis of Force Production by Nascent Kinetochore Fibers. Journal of Cell Biology *113*, 805-815.

Arasada, R., and Pollard, T.D. (2014). Contractile ring stability in S. pombe depends on F-BAR Protein Cdc15p and Bgs1p transport from the Golgi complex. Cell Rep *8*, 1533-1544.

Balasubramanian, M.K., McCollum, D., Chang, L., Wong, K.C.Y., Naqvi, N.I., He, X.W., Sazer, S., and Gould, K.L. (1998). Isolation and characterization of new fission yeast cytokinesis mutants. Genetics *149*, 1265-1275.

Bausch, A.R., Moller, W., and Sackmann, E. (1999). Measurement of local viscoelasticity and forces in living cells by magnetic tweezers. Biophysical journal *76*, 573-579.

Bausch, A.R., Ziemann, F., Boulbitch, A.A., Jacobson, K., and Sackmann, E. (1998). Local measurements of viscoelastic parameters of adherent cell surfaces by magnetic bead microrheometry. Biophysical journal *75*, 2038-2049.

Bezanilla, M., and Pollard, T.D. (2000). Myosin-II tails confer unique functions in Schizosaccharomyces pombe: characterization of a novel myosin-II tail. Mol Biol Cell *11*, 79-91.

Biron, D., Alvarez-Lacalle, E., Tlusty, T., and Moses, E. (2005). Molecular model of the contractile ring. Phys Rev Lett *95*, 098102.

Broersma, S. (1960). Viscous Force Constant for a Closed Cylinder. J Chem Phys *32*, 1632.

Chen, Q., and Pollard, T.D. (2011). Actin filament severing by cofilin is more important for assembly than constriction of the cytokinetic contractile ring. J Cell Biol *195*, 485-498.

Claessens, M.M., Bathe, M., Frey, E., and Bausch, A.R. (2006). Actin-binding proteins sensitively mediate F-actin bundle stiffness. Nat Mater *5*, 748-753.

Clifford, D.M., Wolfe, B.A., Roberts-Galbraith, R.H., McDonald, W.H., Yates, J.R., 3rd, and Gould, K.L. (2008). The Clp1/Cdc14 phosphatase contributes to the robustness of cytokinesis by association with anillin-related Mid1. J Cell Biol *181*, 79-88.

Cortes, J.C., Konomi, M., Martins, I.M., Munoz, J., Moreno, M.B., Osumi, M., Duran, A., and Ribas, J.C. (2007). The (1,3)beta-D-glucan synthase subunit Bgs1p is responsible for the fission yeast primary septum formation. Mol Microbiol *65*, 201-217.

Courtemanche, N., Pollard, T.D., and Chen, Q. (2016). Avoiding artefacts when counting polymerized actin in live cells with LifeAct fused to fluorescent proteins. Nat Cell Biol *18*, 676-683.

Feierbach, B., and Chang, F. (2001). Roles of the fission yeast formin for3p in cell polarity, actin cable formation and symmetric cell division. Current biology : CB *11*, 1656-1665.

Feneberg, W., Westphal, M., and Sackmann, E. (2001). Dictyostelium cells' cytoplasm as an active viscoplastic body. Eur Biophys J Biophy *30*, 284-294.

Hochmuth, R.M. (2000). Micropipette aspiration of living cells. J Biomech *33*, 15-22.

Jochova, J., Rupes, I., and Streiblova, E. (1991). F-actin contractile rings in protoplasts of the yeast Schizosaccharomyces. Cell Biol Int Rep *15*, 607-610.

Kopecka, M. (1975). Isolation of Protoplasts of Fission Yeast Schizosaccharomyces by Trichoderma-Viride and Snail Enzymes. Folia Microbiol *20*, 273-&.

Kovar, D.R., Sirotkin, V., and Lord, M. (2011). Three's company: the fission yeast actin cytoskeleton. Trends Cell Biol *21*, 177-187.

Laplante, C., Berro, J., Karatekin, E., Hernandez-Leyva, A., Lee, R., and Pollard, T.D. (2015). Three Myosins Contribute Uniquely to the Assembly and Constriction of the Fission Yeast Cytokinetic Contractile Ring. Curr Biol *25*, 1955-1965.
Laplante, C., Huang, F., Tebbs, I.R., Bewersdorf, J., and Pollard, T.D. (2016). Molecular organization of cytokinesis nodes and contractile rings by super-resolution fluorescence microscopy of live fission yeast. Proc Natl Acad Sci U S A *113*, E5876-E5885.
Li, Y.J., Christensen, J.R., Homa, K.E., Hocky, G.M., Fok, A., Sees, J.A., Voth, G.A., and Kovar, D.R. (2016). The F-actin bundler alpha-actinin Ain1 is tailored for ring assembly and constriction during cytokinesis in fission yeast. Molecular biology of the cell *27*, 1821-1833.
Lord, M., and Pollard, T.D. (2004). UCS protein Rng3p activates actin filament gliding by fission yeast myosin-II. J Cell Biol *167*, 315-325.
Marks, J., and Hyams, J.S. (1985). Localization of F-Actin through the Cell-Division Cycle of Schizosaccharomyces Pombe. European journal of cell biology *39*, 27-32.
McDonald, N.A., Lind, A.L., Smith, S.E., Li, R., and Gould, K.L. (2017). Nanoscale architecture of the Schizosaccharomyces pombe contractile ring. Elife *6*.
Meyer, R.K., and Aebi, U. (1990). Bundling of actin filaments by alpha-actinin depends on its molecular length. J Cell Biol *110*, 2013-2024.
Mishra, M., Huang, Y., Srivastava, P., Srinivasan, R., Sevugan, M., Shlomovitz, R., Gov, N., Rao, M., and Balasubramanian, M. (2012). Cylindrical cellular geometry ensures fidelity of division site placement in fission yeast. J Cell Sci *125*, 3850-3857.
Ott, A., Magnasco, M., Simon, A., and Libchaber, A. (1993). Measurement of the persistence length of polymerized actin using fluorescence microscopy. Phys Rev E *48*, R1642-R1645.
Pelham, R.J., and Chang, F. (2002). Actin dynamics in the contractile ring during cytokinesis in fission yeast. Natur *419*, 82-86.
Platt, J.C. (1989). Constraint methods for neural networks and computer graphics. PhD Thesis, California Institute of Technology.
Riveline, D., Wiggins, C.H., Goldstein, R.E., and Ott, A. (1997). Elastohydrodynamic study of actin filaments using fluorescence microscopy. Phys Rev E *56*, R1330-R1333.
Shoji, Y., Hamaguchi, M.S., and Hiramoto, Y. (1978). Mechanical-Properties of Endoplasm in Starfish Oocytes. Exp Cell Res *117*, 79-87.
Stachowiak, M.R., Laplante, C., Chin, H.F., Guirao, B., Karatekin, E., Pollard, T.D., and O'Shaughnessy, B. (2014). Mechanism of cytokinetic contractile ring constriction in fission yeast. Dev Cell *29*, 547-561.
Stark, B.C., Sladewski, T.E., Pollard, L.W., and Lord, M. (2010). Tropomyosin and myosin-II cellular levels promote actomyosin ring assembly in fission yeast. Mol Biol Cell *21*, 989-1000.
Takaine, M., Numata, O., and Nakano, K. (2015). An actin-myosin-II interaction is involved in maintaining the contractile ring in fission yeast. J Cell Sci *128*, 2903-2918.
Thiyagarajan, S., Wang, S., and O'Shaughnessy, B. (2017). A node organization in the actomyosin contractile ring generates tension and aids stability. Mol Biol Cell *28*, 3286-3297.
Tseng, Y., Kole, T.P., and Wirtz, D. (2002). Micromechanical mapping of live cells by multiple-particle-tracking microrheology. Biophysical journal *83*, 3162-3176.
Valberg, P.A., and Feldman, H.A. (1987). Magnetic Particle Motions within Living Cells - Measurement of Cytoplasmic Viscosity and Motile Activity. Biophysical journal *52*, 551-561.

Vavylonis, D., Wu, J.Q., Hao, S., O'Shaughnessy, B., and Pollard, T.D. (2008). Assembly mechanism of the contractile ring for cytokinesis by fission yeast. Science *319*, 97-100.
Wang, Y.L. (2005). The mechanism of cortical ingression during early cytokinesis: Thinking beyond the contractile ring hypothesis. Trends in cell biology *15*, 581-588.
Wu, J.Q., and Pollard, T.D. (2005). Counting cytokinesis proteins globally and locally in fission yeast. Sci *310*, 310-314.
Yeung, A., and Evans, E. (1989). Cortical Shell-Liquid Core Model for Passive Flow of Liquid-Like Spherical Cells into Micropipets. Biophysical journal *56*, 139-149.
Yoneda, M., and Dan, K. (1972). Tension at the surface of the dividing sea-urchin egg. J Exp Biol *57*, 575-587.
Yonetani, A., Lustig, R.J., Moseley, J.B., Takeda, T., Goode, B.L., and Chang, F. (2008). Regulation and targeting of the fission yeast formin cdc12p in cytokinesis. Mol Biol Cell *19*, 2208-2219.
